# Modality-Specific and Amodal Language Processing by Single Neurons

**DOI:** 10.1101/2024.11.16.623907

**Authors:** Yair Lakretz, Naama Friedmann, Jean-Rémi King, Emily Mankin, Anthony Rangel, Ariel Tankus, Stanislas Dehaene, Itzhak Fried

## Abstract

According to psycholinguistic theories, during language processing, spoken and written words are first encoded along independent phonological and orthographic dimensions, then enter into modality-independent syntactic and semantic codes. Non-invasive brain imaging has isolated several cortical regions putatively associated with those processing stages, but lacks the resolution to identify the corresponding neural codes. Here, we describe the firing responses of over 1000 neurons, and mesoscale field potentials from over 1400 microwires and 1500 iEEG contacts in 21 awake neurosurgical patients with implanted electrodes during written and spoken sentence comprehension. Using forward modeling of temporal receptive fields, we determined which sensory or abstract dimensions are encoded. We observed a double dissociation between superior temporal neurons sensitive to phonemes and phonological features and previously unreported ventral occipito-temporal neurons sensitive to letters and orthographic features. We also discovered novel neurons, primarily located in middle temporal and inferior frontal areas, which are modality-independent and show responsiveness to higher linguistic features. Overall, these findings show how language processing can be linked to neural dynamics, across multiple brain regions at various resolutions and down to the level of single neurons.

## Introduction

Natural language is considered one of the hallmarks of human cognitive abilities. During the last decades, theories about human language were developed with the fields of modern linguistics and psycholingusitics, providing detailed descriptions of underlying representations and computations. In parallel, in neuroscience, brain imaging and studies of neurosurgical patients has provided a relatively good understanding of where in the brain language is processed, and what are the typical processing times [1, 2]. However, whether and how theories from linguistics can be linked to neural mechanisms in the human brain remains largely unknown.

Psycholinguistic theories suggest that language processing proceeds first along separate modality-specific routes, phonological for spoken words and orthographic for written words. In the case of phonological processing, there is converging evidence suggesting that neural activity in the auditory cortex during speech perception is organized based on dimensions defined by phonological features. Specifically, it was shown that neural activity during the perception of phonemes, the basic units of speech, shows invariance with respect to manner-of-articulation features, which is more dominant compared to place-of-articulation ones [3], also at the single-cell level [4]. Even in infants, brain responses to syllables can be decoded into a series of orthogonal phonetic features [5]. Those findings illustrate how the organization of neural activity in the human brain fits with earlier inferences from linguistic observations [6].

In the case of visual processing, theoretical work based on behavioral evidence from normal and neuropsychological patients suggests that written words are encoded by a list of their abstract, case-invariant letter identities and their approximate positions relative to word beginning and ending [7–9]. However, neural evidence is still disputed. While neuroimaging studies have identified the region and timing in which written words are first encoded, known as the Visual Word Form Area (VWFA) in the left fusiform gyrus, at around 170ms after stimulus onset [10, 11], it remains debatable whether the neural code in the VWFA is compositional, based on letter identities and their approximate positions [12], as suggested by early theoretical work [7–9], or whether neurons are tuned to higher-order combinations of letters - open bigram detectors [13], or local combination detectors [14, 15].

The gap between theory and empirical evidence becomes even larger when concerning higher linguistic information, such as syntax or semantics, where the transition from sensory-dependent to sensory-independent amodal processing is central. At this higher level, there is uncertainty about the relevant brain regions and typical processing times, and scarce evidence about the neural code. For example, for grammatical features, different studies suggest different brain regions and processing times [16–19]. At the semantic level, “concept cells” in the medial temporal lobe fire for specific concepts [20], and are known to respond during presentation of words or sentences, for instance when the names or pronouns refer to their preferred concepts [21, 22], but the neural mechanisms of sentence-level semantic composition remain unclear.

The combination of several factors might explain these current gaps. The most noticeable one is the limited resolution of neuroimaging methods, which remain the main source of data on human language processing. Non-invasive brain imaging has led to tremendous advance in the mapping of the language network [1, 23–25] and typical processing times [26], however it lacks the required spatiotemporal resolution to elucidate the neural code underlying linguistic features. Higher spatiotemporal resolutions, such as measuring single-unit activity, might be indispensable particularly in the case of high-level linguistic information, as suggested by studies on artificial neural language models, which showed that emerging representations of grammatical information in neural networks can be highly sparse, carried by only small fraction of neurons [27, 28]. Another limitation imposed by a lack of spatiotemporal resolution is the necessity to unnaturally boost neural signals, which is often achieved through experimental paradigms that employ linguistic violations, such as syntactic violations. Such violations might generate additional neural activity over and above that related to the target linguistic phenomenon, thus masking fine effects in neural activity related to feature representation.

To address these points, we recorded brain activity from patients (*n* = 21) with pharmacologically intractable epilepsy, implanted with intracranial depth electrodes to identify seizure focus for potential surgical treatment [29, 30]. The data presented is unprecedented in scope including (1) single neuron activity from over 1000 neurons (2) local field potentials from over 1400 microwires (3) intracranial EEG from over 1500 contacts in various areas of the brain. Simultaneous recordings at the level of single units, local field potentials and intracranial EEG provided an opportunity to sample wide brain regions and address the sparseness, as well as spatial and temporal profiles, of linguistic feature coding in the human brain. Patients were presented with stimuli that included minimal pairs of sentences contrasting various linguistic features such as grammatical number or gender (Figure 1A). The sentences were all grammatical and were presented multiple time in both auditory (listening) and visual (fixed-paced reading) form (Figure 1B). The paradigm was designed to separate phonological, orthographic and amodal processing of high-level linguistic information during normal language processing. In the past, the transition from modality-specific to amodal neuronal codes, has been particularly challenging to address. Such transition is evident at the level of the medial temporal lobe where multi-modal invariance of highly selective semantic or conceptual sparse representations has been discovered [20]. Yet the pursuit of amodal processing at the syntactic or sematic levels in other brain regions, especially those traditionally associated with language processing, has been limited by a scarcity of opportunities to record single unit activity in these areas in humans.

**Fig. 1.**
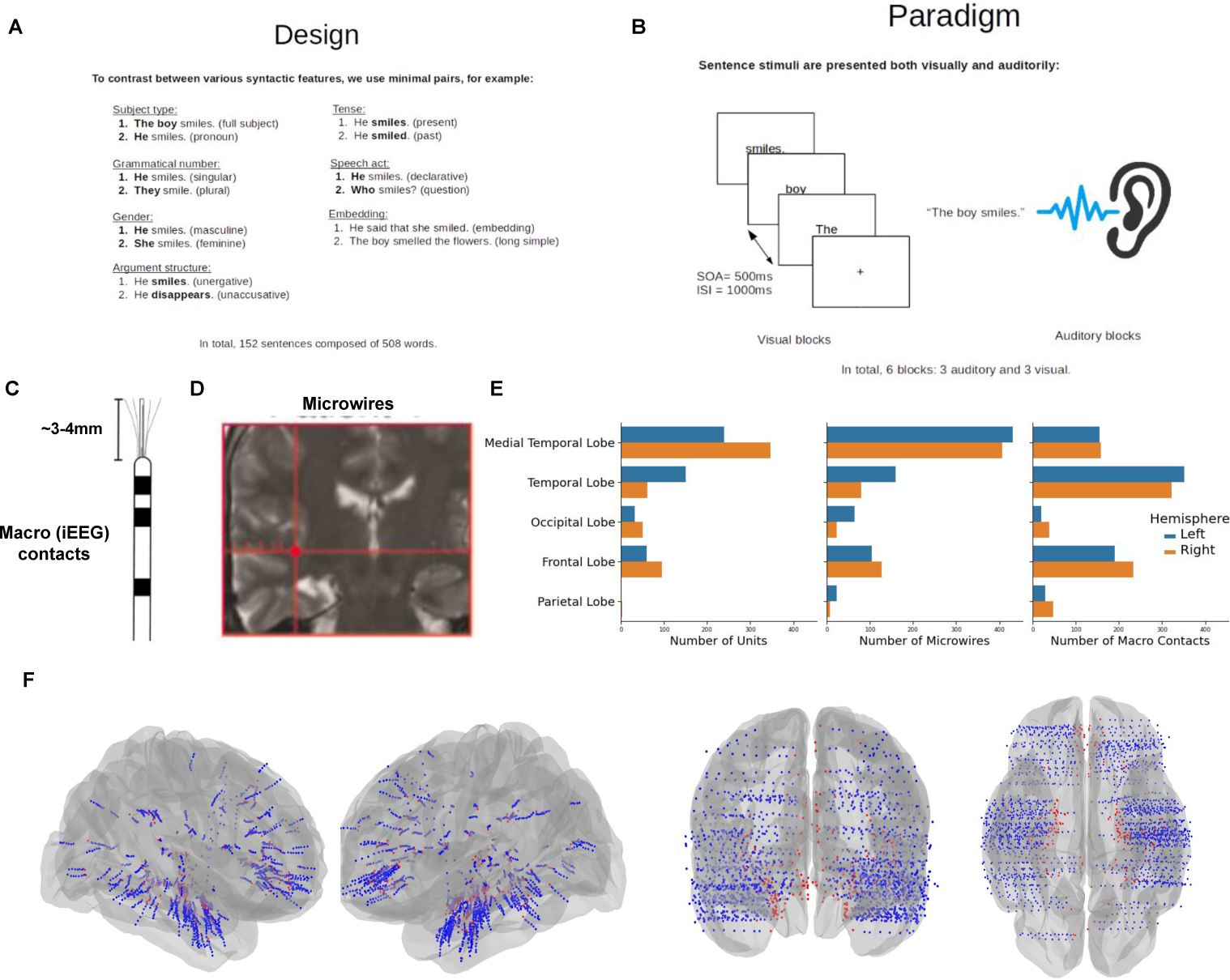
Experimental setup: (A) We recorded single-unit activity from 21 participants, who were presented with 152 sentences, comprising various contrasts along different linguistic dimensions (subject type, grammatical number, etc.). (B) Sentence stimuli were presented to participants both visually, in rapid serial visual presentation (RSVP), and auditorily, via computer speakers. The experiment was composed of six blocks, three from each modality; SOA: Stimulus Onset Asynchrony. ISI: Inter-Stimulus Interval(C) As part of their clinical evaluation, the patients were implanted with Behnke-Fried depth electrodes [31]: the depth electrode contains macro recording contacts along its shaft and 8 microwires protrude from its end, from which single-cell activity can be extracted. (D) an example for a scan of the brain of one of the patients, and the localization of one of the depth electrodes. (E) A summary of the total number of recorded units (left) microwires (middle) and macro recording sites (right). (F) The recording sites aggregated from all the brains of the patients cover main regions from the language network, including ventral temporal cortex, primary and secondary auditory regions, and inferior frontal regions. Blue and red points correspond to macro and micro contacts, respectively.

Using encoding models with flexible temporal receptive fields, we identified single cells in visual and auditory regions that showed high selectivity to only a single modality. Furthermore, some of these single cells showed high selectivity to modality-specific sub-word features such as a letter or a phoneme type. In higher-level regions, particularly in the MTG and IFG, we found modality-independent cells that were active in response to linguistic input from both modalities, some of which showed modality-independent preference for linguistic features such as questions over declarative sentences.

Together, these findings suggest that low-level linguistic information, such as phonology and orthography, are encoded in a local and modality-specific way, whereas higher-level features are encoded in an amodal way, and possibly in a more distributed way. This suggests a local-to-distributed encoding scheme along the processing hierarchy in the human language network.

## Results

Neural recording in patients were conducted with intracranial Behnke-Fried electrodes [31]. Behnke-Fried electrodes have microwires protruding from the tip of the shaft (Figure 1C), from which spiking activity can be extracted (*Methods*). From the microwires and macro contacts along the shaft, we also extracted and computed local field potentials and broadband-gamma activity (BGA), which, together, provide three types of resolutions into neural activity – spiking activity, micro and macro neural-population activity. Figure 1D shows an example of an electrode implanted in one of the patients, illustrating the locations of the macro contacts (small dots) and the target of the microwires (large red dot). Across all patients, based on clinical considerations, most of the microwires were located in the Medial Temporal Lobe (MTL), followed by lateral temporal, frontal and occipital regions (Figure 1E). Macro contacts were mostly present in the temporal and frontal lobes, given the implantation procedure. Figure 1F shows the total coverage from the 21 patients, for micro (red) and macro (blue) recording sites.

### Single-Cell and Neural-Population Responses to Sentences Presented in either the Visual or Auditory Modalities

#### Modality Selectivity of Single Cells in the Fusiform and Superior Temporal Gyri

The Fusiform Gyrus (FSG) and the Superior Temporal Gyrus (STG) are brain regions respectively associated with early orthographic and phonological processing of linguistic input, as shown by numerous neuroimaging and intracranial studies [11, 32–35]. However, their single-neuron activity during language processing has been only scarcely described [4, 36], and spiking activity in response to stimuli from the opposite modality (e.g., visual for STG) has not been previously explored.

Figure 2A shows spiking activity from an example neuron from the fusiform gyrus during the processing of all sentences from the experiment, where trials are sorted based on the number of words in the sentence (2 to 5 words). During visual blocks (top), spiking activity was strongly elicited ∼ 170*ms* after sentence onset (vertical dashed line), matching that of the N170 component of local fields associated with orthographic processing in the fusiform gyrus [11]. Spiking activity returned to baseline after a short period of a few tens of milliseconds, and was elicited again by the next word, with a temporal delay corresponding to the spacing between words in the paradigm (SOA=500ms), until the end of the sentence (vertical red lines). In contrast, during auditory blocks (bottom), the spiking activity of this fusiform neuron remained at baseline throughout the entire auditory presentation of sentences. Together, this shows the high selectivity of the fusiform neuron to the orthographic-visual modality, which was also observed in several other neurons in the dataset (Figure B2).

**Fig. 2.**
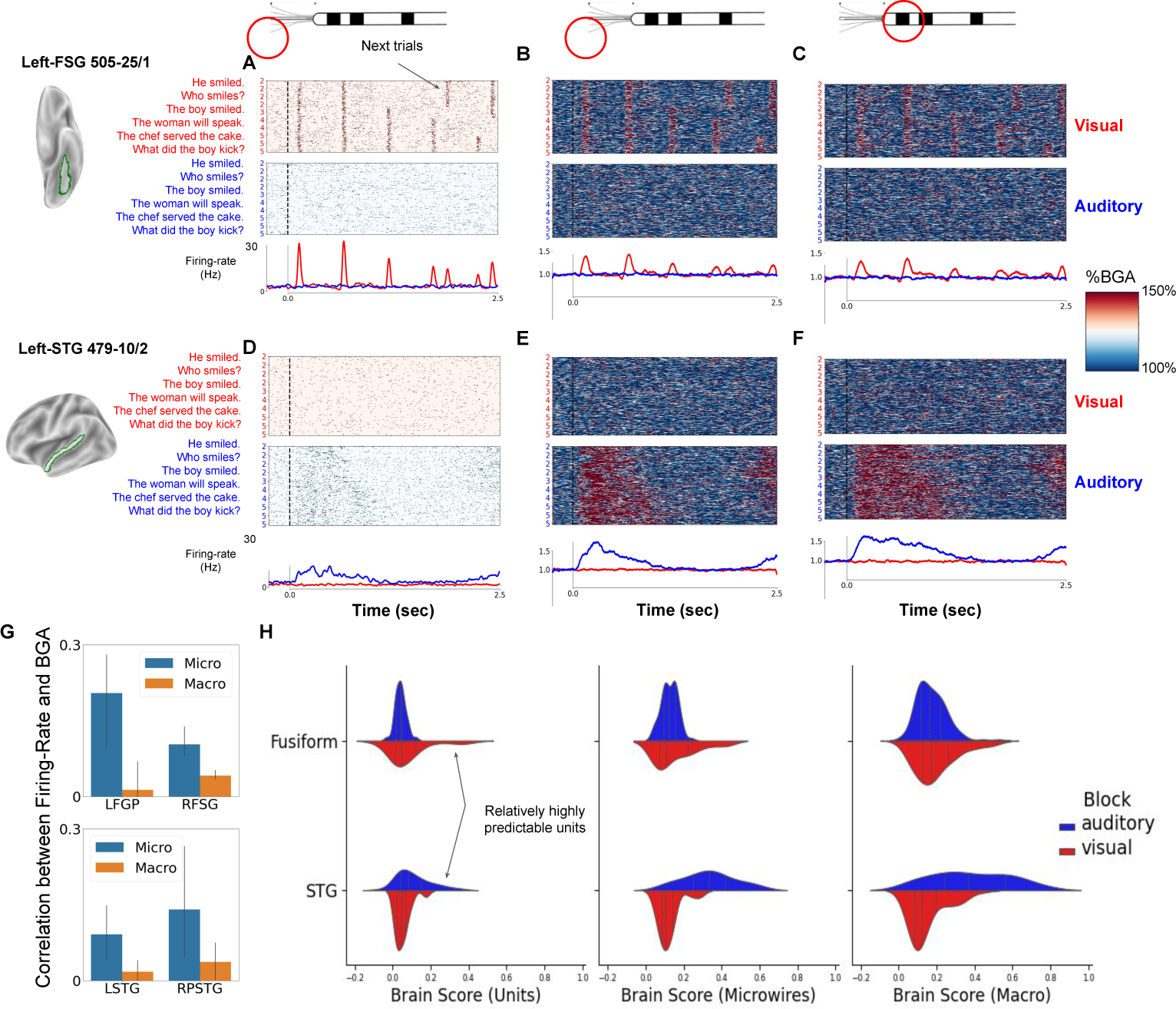
From single cells to large neuronal populations: (A) An illustration of spiking activity of a neuron from the left fusiform gyrus of one of the patients during the processing of all sentence stimuli. (B) The corresponding broadband-gamma activity (BGA) extracted from the *same* microwire as the neuron. (C) BGA extracted from the nearest macro contact. (D-F) Same for an example single neuron and the corresponding micro- and macro-BGA from the superior temporal gyrus (STG) of another patient. In all panels A-F, trials are sorted based on the length of the sentence (2-5 words). Sentence onset is marked by a vertical dashed line. The estimated time-resolved firing rate and the median BGA are shown at the bottom of each panel. In each panel, top and bottom panels show activity during the visual and auditory blocks, with red and blue backgrounds, respectively. In the fusiform gyrus, neural responses are selective to visual compared to auditory stimuli. For all the three resolutions (single-cell to micro- and macro-population activity), neural responses are highly selective to visual compared to auditory stimuli. In contrast to the fusiform gyrus, neural activity at the STG is highly selective to auditory compared to visual stimuli, also at the single-cell level. (G) Mean Pearson correlation between firing-rate and high-gamma activity for FSG (top) and STG (bottom) units during visual and auditory blocks, respectively. Blue bars correspond to correlation between spiking activity and high-gamma extracted from the same microwire. Orange bars correspond to correlation between spiking activity and high-gamma extracted from the nearest macro contact. (H) Left: Brain-Score distributions for all recorded single units from the fusiform gyrus (top) and STG (bottom), computed separately for the visual (red) and auditory (blue) blocks. Middle: same for microwires; Right: Same for macro contacts.

An opposite modality selectivity was observed in STG neurons. Figure 2D shows spiking activity of an example neuron from the STG of one of the patients. During auditory blocks (bottom), spiking activity was strongly elicited following a relatively short interval (∼ 50*ms*) after sentence onset (vertical dashed line). Spiking activity continued until the end of the sentence, following the duration of the continuous speech that was presented to the patient during the auditory blocks. In contrast, during visual blocks, the STG neuron was mostly silent throughout the entire visual presentations of the stimuli, without any rhythmic firing at the 500 ms period with which the visual stimuli were presented. This shows the high selectivity of the STG neuron to the auditory modality, which was also observed in many other STG neurons in the dataset, as well as in Heschel’s gyri (Figures B3&B4).

#### Modality Selectivity of Small Neuronal Populations in the Fusiform and Superior Temporal Gyrus

From the same microwires, we extracted broadband-gamma activity (BGA), which represents local neuronal-population activity in the vicinity of the above two single cells. Figures 2B&E show BGA during the processing of the sentences, in both visual and auditory blocks. Figures 2C&F further show BGA extracted from the nearest macro contact (with bi-polar referencing; Methods). For micro BGA in the fusiform gyrus, local population activity was high during visual blocks, but absent during auditory blocks. In contrast, in the STG, population activity was high during auditory blocks, but absent visual blocks. This shows that the modality selectivity observed at the single-cell level occurs also at the local population level. Overall, Macro BGA had a similar response duration, for both block types. However, a careful examination of the response profiles reveals larger discrepancy with spiking activity, compared to micro BGA, as described next.

#### Spiking Activity Correlates Stronger with Micro compared to Macro High-Gamma Activity

We next quantified the similarity between single-cell spiking activity and that of micro and macro BGA, which correspond to larger neuronal population activity. For this, for all responsive (Methods) FSG and STG units (Figures B2-B4), we computed the Pearson correlation between the firing rate of the neurons and either micro or macro BGA. Figure 2G shows that spiking activity correlates with micro BGA (blue bars), for both FSG and STG units (*p* − *value <<* 0.05), but to a much lesser extent with macro BGA (orange bars).

#### Predictability of Single-Cell Activity in the FSG and STG by Linguistic Features

We next studied whether linguistic features structure neural activity of single-cells. For this, we first estimated which linguistic information (e.g., orthographic, phonological, syntactic) best accounts for observed evoked neural activity, focusing on six possible feature groups (Table A2), using encoding (forward) modelling of temporal receptive fields (TRFs). TRFs model neural activity at a given time point as a linear combination of features occurring at preceding time points (see Methods). We further estimated the causal role of each linguistic feature in predicting neural activity, beyond mere correlation with it, using ablation methods applied to the TRFs. Specifically, for each single cell and recording contact, we compared the performance of the full TRF model and that of an ‘ablated’ model, in which we removed features of interest (see Methods). We refer to the resulting drop in model performance as *feature importance* when only a single feature was removed from the model, and *feature-group importance* when a group of features was removed (e.g., removing all syntactic features). In the case of visual presentation, we included orthographic but not phonological features in the TRFs, and vice versa in the case of auditory presentation. For model evaluation, we defined *Brain-Score* as the Spearman correlation between predicted and true neural activity in the unseen set. Since TRFs are computed for each modality separately, each unit and electrode is described by two Brain-Scores, one for the visual and one for the auditory modality. In all cases, TRFs and feature-importance values were evaluated on unseen data in a cross-validation manner (*Methods*).

Figure 2H shows the Brain-Score distributions for the three levels of recording resolutions - spiking activity (left), micro (middle) and macro BGA (right), for the fusiform (top) and STG (bottom) and for both the visual (red) and auditory (blue) modality. In the fusiform gyrus, for all recording types, Brain-Score distribution had a longer tail for visual compared to auditory blocks, indicating that many units and sites responded in a highly predictable manner only to visual stimuli. Neural activity in the fusiform is thus more predictable by linguistic features during reading compared to listening. Conversely, neural activity in the STG was more predictable by linguistic features during auditory blocks, having longer-tail distributions for all three recording types. The maximal Brain-Score for single cells in the FSG and STG was *Brain* − *Score* = 0.42, 0.64, respectively, suggesting that spiking activity in these regions is relatively highly predictable by linguistic information. Maximal Brain-Scores for micro and macro BGA were even higher, both above 0.8.

### Single-Cell Encoding of Orthographic Information in the Fusiform Gyrus

#### Selective Single-Cell Responses to Letters

We next studied single neurons in the fusiform gyrus with relatively high Brain-Score during the visual block. We analyzed which linguistic-feature groups best predict their spiking activity, and whether their response is selective to specific features.

Figure 3A shows feature-group importance of all five groups for an example neuron from the right FSG, computed based on spiking activity during visual blocks only (Brain-Scores from the auditory TRF were not statistically significant). Results show that the importance of the orthographic-features group was significantly higher compared to all other groups, and the only one that was significantly above zero, which suggests that this cell mostly or solely encodes orthographic information.

**Fig. 3.**
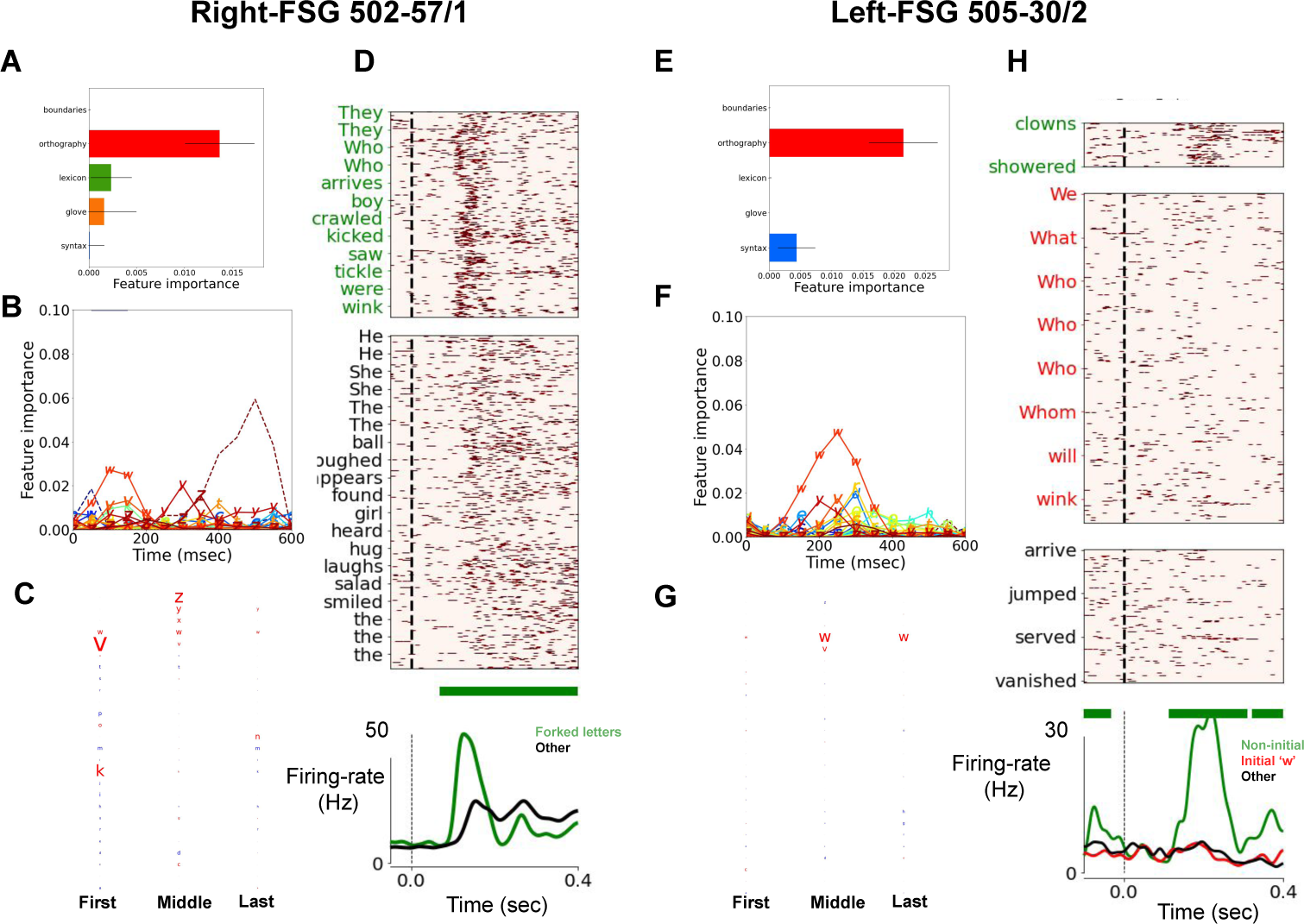
Selective encoding of orthographic features in the fusiform cortex: To study the information encoded in firing patterns of single neurons, we fitted firing rates to *diagnostic encoding models* which contained features from only the most dominant feature group in the visual or auditory-block model. (A-D) An in-depth analysis of an example single neuron from the fusiform gyrus. (A) The most dominant feature-group in the visual-block model was for orthography. Feature importance was measured as the difference in model performance with vs. without each feature group. Bars correspond to the mean significant feature importance within 0-600 msec (FDR corrected). (B) Given the results of the visual-block model, the diagnostic model contained features only from the orthography group - letter identities and word length. The letter features that achieved highest importance score were: ‘k’, ‘v’, ‘w’, ‘y’ and ‘z’. Note that all these letters contain a V-shape feature in various orientations. (C) To study the effect of letter position on spiking activity, we trained a second diagnostic model, which contained features that couple letter identity (26 letters) and letter position (3 levels, depending on whether the letter appeared at the beginning, middle, or end of the word). The resulting weights of the model features are plotted for each of the three positions, with font size corresponding to the size of the weight. Red and blue represent positive and negative values, respectively. (D) An illustration of the spiking activity of this neuron, grouped based on whether the word contains (top panel) or not (bottom) a letter with a V-shape feature. Word onset is marked with a vertical dashed line. The bottom panel shows the mean firing rate for each group. (E-H) Same analyses for another example neuron, which shows high selectivity to a single letter, with increased firing rate in response to words that contain the letter ‘w’, only if it appeared at the middle or end positions.

For this cell, we next studied the importance of each feature in the orthographic-features group. Figure 3B shows the time-resolved feature importance for all 26 letter identities in the English alphabet, which were included in the encoding TRF model. Results showed that only the occurrence of a specific subset of letters (’k’, ‘v’, ‘y’ and ‘z’) elicited a significant change in spiking activity. Note that the underlying commonality of these letters is that they all contain a V shape in a given orientation. These letters are known in cognitive science as ‘forked letters’, and the ‘forking’ feature was previously suggested, purely on the basis of behavioral studies, as an important discriminative feature of letters [37].

We next studied whether spiking activity of this cell depends not only on the occurrence of a forked letter in the word, but also on the position in which it occurs. For this, we fitted another TRF model, in which letter identities were coupled to three possible positions (first, middle and last position in the word). Figure 3C shows feature importance of each letter-in-position feature. It shows that ‘k’ and ‘v’ elicited spiking activity when in the first position, and ‘y’ and ‘z’ in the middle one. This finding is consistent with the hypothesis that written words are encoded orthographically by assemblies of neurons sensitive to specific letters at an approximate position, a neural code which was previously proposed only on the basis of behavioral and simulation studies [8, 9, 12, 38, 39]. The lateralization of this neuron to the right FSG is compatible with its response to contralateral letters.

Finally, to illustrate the results from the TRFs, Figure 3D contrasts spiking activity in response to words that contain or do not contain forked letters. It shows that the mean firing rate to words that contain forked letters is significantly greater compared to those that do not contain forked letters (temporal cluster-based permutation, p-value ¡ 0.05; green horizontal strip). Comparing with micro BGA extracted from the same microwire, here too, we found correspondence between spiking and local high-gamma activity. Figure C5A shows a significant separation between micro-BGA during the processing of words that contain forked letters and those that do not (temporal cluster-based permutation, *p* − *value <* 0.05; green horizontal strip). In contrast, such separation was not observed for macro BGA Figure C5B).

Together, this shows how letter features, such as forking, described by theoretical work purely based on behavioral data can be linked to small neuronal populations, and down to the single-cell level.

A similar analysis of another single cell from the Fusiform Gyrus revealed an even higher selectivity compared to that described in the previous section. This cell was highly selective to words that contained a specific letter, and only when it occurred at specific positions in the word. Figure 3E shows the feature-group importance for this unit, showing that spiking activity was solely selective to orthographic information. Figure 3F further shows the time-resolved feature importance of all letters, where spiking activity around 150-400ms was mostly explained by the occurrence of the letter ‘w’ within the current word. Finally, Figure 3G shows letter-in-position feature importance, suggesting that spiking activity is selective to the occurrence of ‘w’ in middle and last positions (e.g., ‘flowers’), but not when it occurs at the beginning of the word (e.g., ‘will’), in fitting with the left-hemispheric localization of this neuron. Finally, Figure 3H illustrates this neuron’s selectivity by grouping words into whether they (1) contain ‘w’ in middle or last position, (2) ‘w’ at the beginning of the word, and (3) do not contain ‘w’. The mean firing rate rises following the stimulus word only in the first case, but not the other two. Note that the case of the letter ‘w’ (lower vs. upper) cannot explain the difference between (1) and (2) – firing rate was at baseline also when ‘w’ appeared at the beginning of words in non-initial positions of the sentence (e.g., ‘winking’). Comparing with micro BGA extracted from the same microwire, as in the previous case, we found correspondence between spiking and local high-gamma activity. Figure C5C shows a significant separation between micro-BGA during the processing of words that contain ‘w’ in middle positions compared to those that do not (temporal cluster-based permutation, p-value ¡ 0.05; green horizontal strip). Such separation was found to a much lesser extent, yet significant during a shorter duration, for macro BGA Figure C5D).

Taken together, these analyses show that single cells in the Fusiform Gyrus can be highly selective to the visual modality, to specific orthographic features (forking) or to letter identity in specific positions in a word. In total, we have recorded neural activity from neurons from the FSG that were responsive to the stimuli. Figure B2 shows raster plots from seven neurons with strongest responses and Brain-Score (Methods), as well as their feature-importance profiles, showing high selectivity to the group of orthographic features in all cases.

#### Single-Cell and Population Encoding of Word Length

We next studied the neural encoding of the word-length feature (number of letters), which was included in the TRF analysis as another non-linguistic covariate. The feature importance of word length in the TRFs was significant for a large number of units and electrode contacts. We first visualized the effect of word length on neural activity by projecting the activity from all responsive FSG neurons onto the two main principal components (PCs) of the data. Figure 4A visualizes all words in the dataset that were presented more than 10 times during the experiment, along the two first two PCs of spiking activity (*Explained* − *V ariance_P C_*_1_ = 8.5%, *Explained* − *V ariance_P C_*_2_ = 7.1%). Words are colored based on their length, with cool and hot colors for short and long words, respectively. Results show that, qualitatively, the first PC of spiking activity corresponds to word length, with longer words (red) having larger values along PC1. To quantify this, we computed the Pearson correlation between the first PC and word length, which was significant *r* = 0.31 (*p* − *value <* 10^−3^). Given the previous results on forking letters, we further observed that, qualitatively, words with forked letters tend to span larger values of PC2. We computed the Pearson correlation between the second PC and the number of forked letters in the word, which, indeed, was found significant *r* = 0.45 (*p* − *value <* 10^−3^). In contrast, correlation between PC1 and number of forked letters was not significant, as well that between PC2 and word length.

**Fig. 4.**
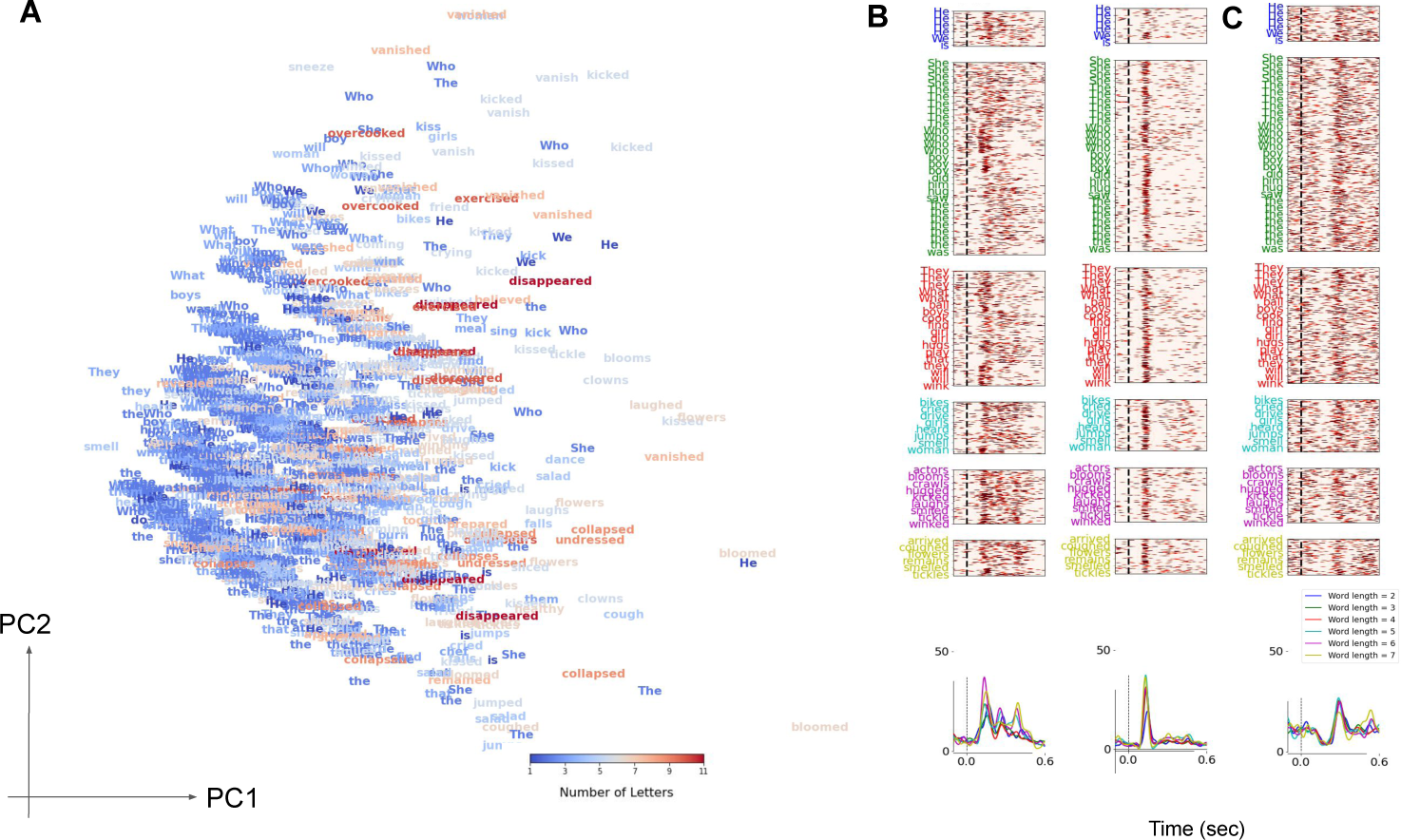
Neural Encoding of Word Length: (A) Principal Component Analysis (PCA) of spiking activity from the 5 fusiform neurons with the highest Brain-Score. The neural response of each neuron was represented along 5 consecutive 100ms time bins between 0.1 and 0.6 seconds after word onset. The population response to different words is projected onto the two main PCs. Words are colored by their length. (B) Raster plots from two example neurons from the FSG with significant feature-importance of word length. Different raster panels correspond to different word lengths (y-label colors). Bottom panel shows the mean firing rate across trials, split into different word lengths (same color code as y-labels), which increases as a function of word length. (C) Same as previous panel but for a neuron from the FSG whose activity is not sensitive to word length.

Figure 4B further illustrates the effect of word length on neural activity at the single-cell level. It shows raster plots from two units with significant feature importance for word length, grouped by words with different lengths (different colors). The mean firing rate (bottom panels) of these neurons increased with respect to word length, in contrast to other units, as illustrated by Figure 4C.

Finally, comparing to micro BGA recorded from the same microwires, the same analysis found significant, albeit weak, correlation between the two first PCs and word length (*r* = 0.11, *p* − *value <* 10^−3^) and forking (*r* = 0.07, *p* − *value <* 0.002), respectively (Figure F12). Together, these results suggest the dominance of word-length effect in population activity in the fusiform gyrus.

### Single-Cell Encoding of Phonological Information

#### Selective Single-Cell Responses to Consonants in the Auditory Cortex

An analysis of spiking activity of single cells in the auditory cortex revealed a similar selectivity to phonological information. Figure 5A shows feature-group importance for a single cell from the left Heschel Gyrus (l-HG). Spiking activity of this cell is highly selective to the phonological feature group, but to less extent to higher-level information. Figure 5B shows the time-resolved feature importance of all phonemes, suggesting that spiking activity around 50-200ms is mostly explained by the occurrence of the phoneme */SH/* (as in ‘she’; using the ARPAbet code, 40). Finally, Figure 5C shows letter-in-position feature importance, suggesting that spiking activity is selective to the occurrence of */SH/* and */JH/*, both are hushing-sibilant phonemes. Figure 5D illustrates this with raster plots for example phonemes, including other similar sibilants such as */S/* and */Z/*.

**Fig. 5.**
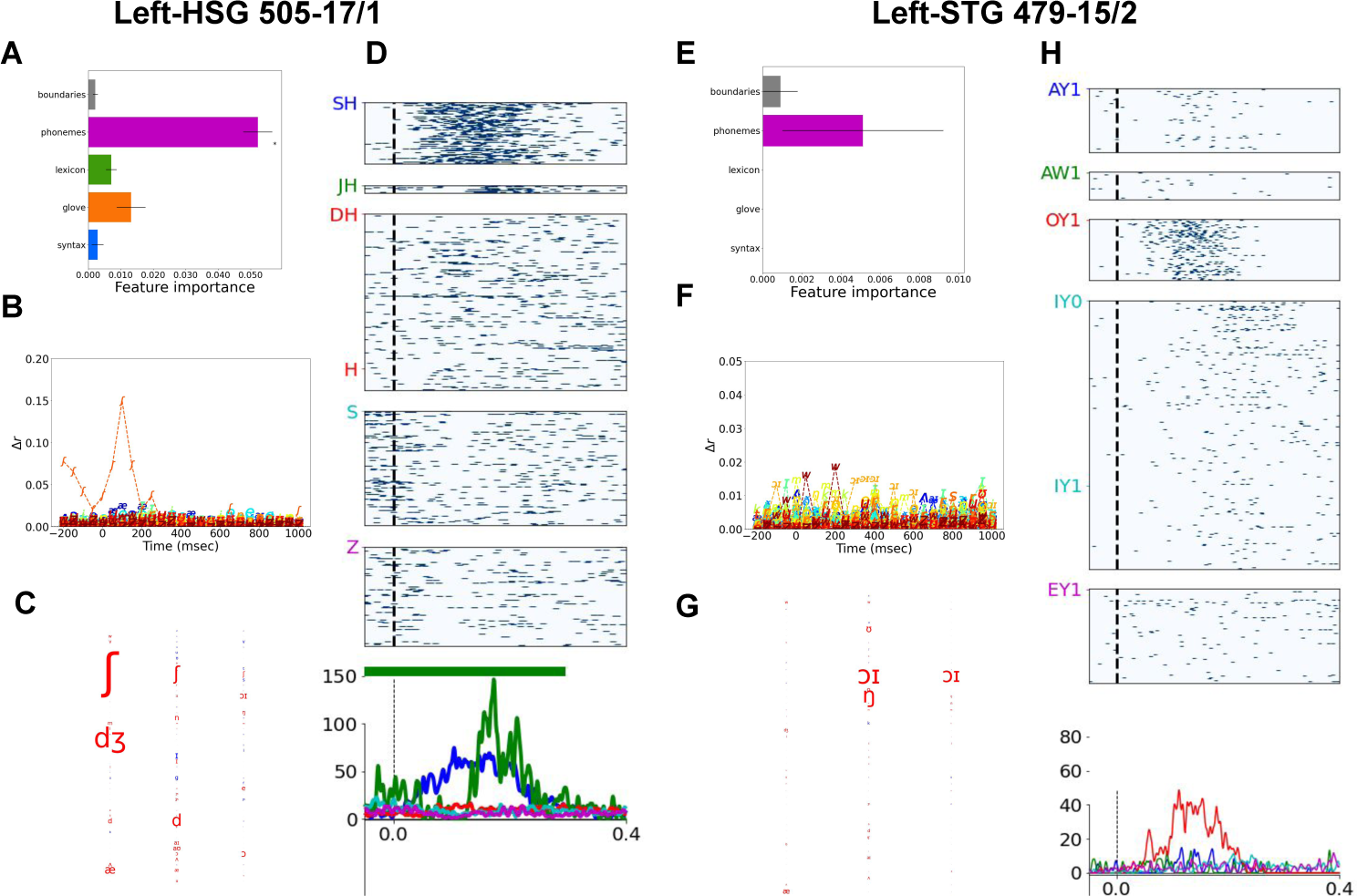
Selective encoding of phonological information during auditory processing: (A-D): An in-depth analysis of an example single neuron from the left Heschel Gyrus. (A) Feature-group importance for the five groups in the temporal receptive field (TRF). The feature group that had largest effect on spiking activity was that of phonological features. (B) Time-resolved feature importance for all phones. The phone that had the largest effect on spiking activity was the phone */SH/*. (C) Phone-in-position feature importance, showing the effect of the phone */SH/* and */JH/* on spiking activity. (D) Raster plots for several example phones, illustrating the high selectivity of this cell to the phone */SH/*. Vertical dashed black lines represent phoneme onset. Horizontal vertical line represents period with a significant difference among conditions (a temporal cluster-based permutation test, *p − value <* 0.05) (E-H) Same analyses for another example cell, which shows high selectivity to a single vocalic diphthong.

Comparing with micro BGA extracted from the same microwire, in contrast to results from the FSG, we did not find correspondence between spiking and micro high-gamma activity. Figure C6A shows that there’s no significant separation between micro-BGA during the processing of words that contain hushing sibilants compared to those that do not; nor for macro BGA (Figure C6B)

#### Selective Single-Cell Responses to Vowels in the Auditory Cortex

A similar analysis of another single cell from the auditory cortex, with a relatively high Brain-Score, revealed a similar selectivity, but this time to a specific diphthong rather than a consonant. Figure 5E shows the feature-group importance of the five groups for a single cell from the left Superior Temporal Gyrus (l-STG). In this case too, spiking activity of this cell is highly selective to phonological but not to higher-level information. Figure 5F further shows the time-resolved feature importance of all phonemes, suggesting sensitivity to */OY/* (as in ‘boy’) during the time interval of 250-400ms. Finally, Figure 5G shows sound-in-position feature importance, suggesting that spiking activity is selective to the occurrence of */OY/*. Since the diphthong */OY/* did not occur in first position in our dataset, we cannot conclude whether this cell is selective to its occurrence in middle and last position, as suggested by the TRF. Figure 5H illustrates the TRF results with raster plots for various diphthongs.

Comparing with micro BGA extracted from the same microwire, we found correspondence between spiking and micro high-gamma activity. Figure C6C shows that there’s significant separation between micro-BGA during the processing of words that contain */OY/* diphthong and those that do not, although to a lesser extent compared to spiking activity. Such separation was not observed for macro BGA (Figure C6D).

### Amodal single-cell activity at the Middle Temporal Gyrus and Inferior Fronal Gyrus

We next turned to high-level processing, putatively involved in abstract amodal language encoding. Modality preference of single cells and recording contacts was quantified comparing Brain-Scores from the two input modalities. Figures 6A shows the difference in Brain-Score between the two modalities, for all recording sites (single cells, micro and macro electrodes. In accordance with the qualitative observations above (Figure 2), visual and auditory brain regions show significant preference to a single modality (red and blue colors, respectively), whereas higher-level regions in the language network, including the posterior temporal sulcus, middle temporal gyrus and the inferior frontal gyrus, show amodal processing (in green). Figure 6B further contrasts the two Brain-Scores for different brain regions - (1) visual areas (red hues), including the fusiform gyrus, (2) auditory areas (blue hues), including the superior temporal gyrus, and (3) amodal regions (in green), including part of Broca area (Brodmann 45). Finally, we illustrate neural activity in five identified amodal single neurons, from the right anterior insula and from the middle temporal gyrus from two of our patients (Figure 6C). In all cases, an evoked response occurred after the presentation of the stimuli in both modalities, in contrast to modality-specific responses of single cells from earlier processing regions (Figure 2, Figure B2-B4).

**Fig. 6.**
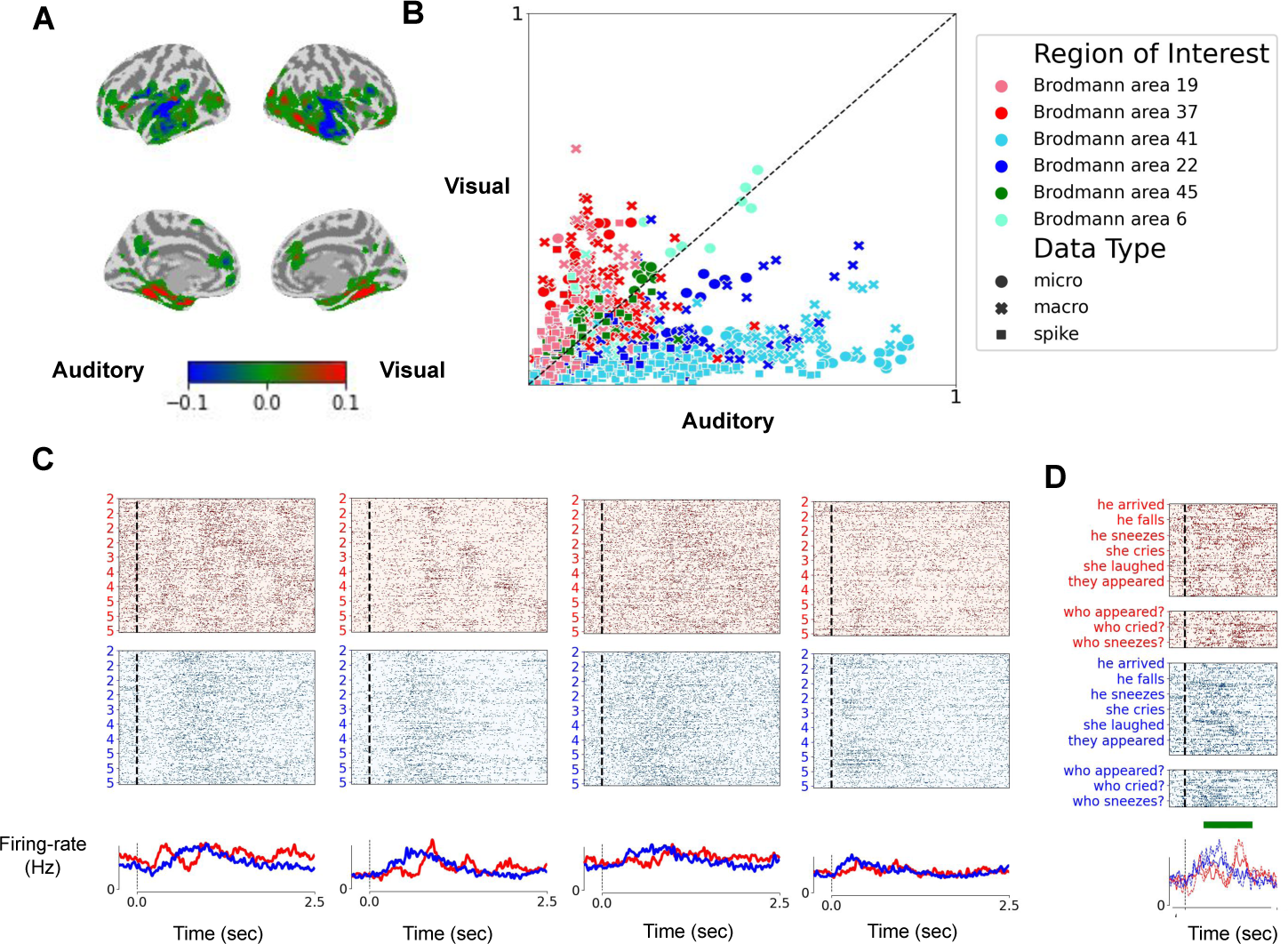
Modality-specific and Amodal Activity across the language network: To quantify modality specificity of all neural responses, for each neuron and for each recording electrode, we trained two encoding models, one for each of the two modalities - a *visual-block* and *auditory-block* models. The models contained features from various linguistic aspects: The visual-block model contained orthographic, lexical, semantic and syntactic features, as well positional features, such as sentence onset or word position. The auditory-block model contained the same features but ortho-graphic features were replaced with phonological ones. We evaluated the visual- and auditory-block models by testing their performance in predicting unseen neural data, in a cross-validation (CV) procedure. We define the *Brain-Score* as the mean performance of the models across all CV splits. (A) Brain-Score differences between the auditory and visual blocks. Red and blue shades correspond to visual and auditory specificity, respectively. Green corresponds to amodal regions. (B) A scatter plot illustrating Brain-Score for auditory vs. visual block models. Each dot corresponds to a single-cell (squares), a microwire (circles) or a macro contact (X marks). Colors correspond to selected regions of interests. Dots on the lower/upper triangle correspond to recording sites with preference to auditory/visual stimuli. Dots on the diagonal corresponds to amodal recording sites. (C) Single-cell activity of several amodal cells from the MTG and right IFG. (D) Spiking activity of an MTG amodal cell, in responses to two-word sentences, either questions (dashed lines) or declaratives (continuous).

A TRF analysis of the activity of the amodal neurons reveals a complex encoding profile, with several feature groups showing a significant importance. Figure G13 contrasts the FIs from the two blocks (visual and auditory), showing that FI values often depend on input modality. However, for all cells, the syntax feature group reaches a relatively high feature importance in visual blocks. An analysis of one of the amodal cells in the MTG (Figure 6D) shows a slight, yet statistically significant, preference to questions versus declarative sentences, with firing rates that are significantly stronger for the former in both modalities (temporal cluster-based permutation test; *p* − *value <* 0.05; green strip). However, across all units in our dataset, we did not observe strong selectivity to only a single high-level linguistic feature. This stands in contrast to lower-level modality-specific processing of orthography and phonology.

## Discussion

We present a new dataset and analyses of neural activity of single cells and large neural populations in 21 neurosurgical patients during the processing of sentences. The new dataset suggests an unprecedented examination of linguistic features across various brain regions at various resolutions down to the level of single neurons, and processed by visual and auditory modalities, during reading and listening, respectively.

Our data demonstrates a double dissociation between phonological and orthographic processing by single neurons in fusiform gyrus and superior temporal gyrus. This was enabled by our recording methodology enabling single neuron recordings in these two regions, coupled with bimodal presentation of stimuli to the same patient. A stark demonstration of this double dissociation is demonstrated in patient 502 with neurons in HSG (Figure 5) recorded simultaneously with neurons in FSG (Figure 3). During reading, we found specific neurons in the fusiform gyrus that were highly selective to orthographic information compared to other linguistic features. These neurons are located around the fusiform gyrus, where the visual word form area (VWFA) resides [41], a brain area that was shown in neuroimaging studies to encode written-word information [see, 11, for a review]. Our data provide a first glimpse into how neurons in the vicinity of the VWFA encode orthographic information. The activity of the neurons in the fusiform gyrus showed high selectivity to written compared to heard words. Neural activity after the visual presentation of a word spikes at around 170ms after word onset, with a mean firing rate that goes up to 30Hz, whereas after auditory presentations of the same words, neural activity of these neuron remains at baseline close to 0Hz. Previous work has identified modality preference during language processing [41, 42], however, to our knowledge, this is the first demonstration of such modality preference during reading at the single-cell level in humans (for similar work in monkeys exposed to written words, but obviously unable to read them, see 43).

Several studies have shown that the role of the VWFA may extend beyond mere bottom-up visual word recognition, also encompassing aspects of speech perception. While the VWFA is typically inactive during passive listening to spoken words [44], it can be activated in a top-down manner under certain circumstances [45]. This activation is more likely to occur during complex speech tasks, such as lexical decision or rhyming, and in individuals with strong literacy skills [14, 46, 47]. Interestingly, more recently it was shown that in individuals with enhanced connectivity between high-level cortical regions and the VWFA, VWFA activation can further increase during speech perception, i.e., when linguistic input was presented in the auditory modality [48]. This suggests that while the primary function of the VWFA is orthographic processing, it can also be recruited to support speech perception, suggesting a more interconnected language processing system with possible top-down influences from phonological and higher-level language areas. In contrast, the strong modality selectivity observed in our data suggests that during speech perception, the firing of many units in the fusiform are not subject to such top-down activations during speech perception. Similarly, in the converse direction, we found a large number of units in Heschl’s gyrus and STG that are silent during reading (Figures B2-B4). The activity of these units is poorly predicted by linguistic features during visual blocks (i.e., low Brain-Score). This raises questions for future research, to further disentangle the interplay between bottom-up processing, as frequently observed in our data at the single-cell level, and top-down processing. One possibility is that bottom-up processing occurs at deeper cortical layers, whereas top-down processing during reading and speech comprehension occurs in possibly more superficial layers [49–51]. While we cannot address layer-of-origin in the present data set, this limitation could be overcome with novel recording techniques, such as neuropixel probes [36].

Our dataset also provides a unique opportunity to compare single-cell activity and neural activity at two scales, both at the level of the microwire and that of macro contact. For both the STG and the FSG, we found overall stronger correlation between spiking activity and micro-BGA, extracted from the same microwires, compared to correlations between spiking activity and macro-BGA, extracted from the nearest macro contacts (Figure 2B). The relatively strong correlation with micro-BGA suggests that information about the inputs may be encoded redundantly at a local level, across many neurons in the vicinity of the microwire. That is, to have a measurable impact on high-gamma activity, as we found, neural populations at the vicinity of the microwire need to encode information in a sufficiently redundant manner. In contrast, the correlation between spiking activity and macro-BGA decreased significantly, indicating that redundancy might diminish drastically over a distance of 2-3 millimeters.

Classic cognitive work on letter similarity, based on behavioral data, has identified distinct groups of letters, separated into clusters [37], such as ‘forked letters’, a cluster that appears to selectively engage the fusiform gyrus neuron shown in Figure 3. Behavioral and fMRI data suggest a compositional code for orthography, whereby letter identities and letter positions are jointly encoded [8, 12]. Simulations of how convolutional neural networks for visual recognition can be “recycled” to recognize written words, also suggested a similar letter-by-position code [38, 39]. However, whether such a code actually exists at the single neuron level has been an open question. Our findings provide the first direct evidence in support of this theory. Figure 3B shows a neuron from the left fusiform gyrus neuron, which was found to be selective to a single letter ‘w’, but only when the letter appeared in middle or last positions of the word. Such a coupling between letter position and letter identity, although needed to be described in more such cases, is in accordance with a compositional code for orthography

The increased selectivity to more complex features, from sub-letter features (e.g., ‘forking’) to single letters, is in accordance with hierarchical theories of information processing across the visual cortex. The observed selectivity of single neurons is compatible with the hypothesis that ventral occipitotemporal cortex responses to abstract letter identities are based on a hierarchy of earlier visual stages, as was previously shown with fMRI [10] and sEEG data [35], with initial bilateral processing of contralateral letters followed by a progressive convergence towards left-hemispheric cortical patches forming the VWFA [41, 52]. The units that we report here, including a right fusiform neuron sensitive to left-hemifield letters and a left fusiform neuron sensitive to non-initial letters, are likely to lie within this initial contralateral path, although they might reflect a remnant contralateral bias within the VWFA itself, which has been reported with fMRI [53].

During auditory processing of speech, modality-specific selectivity at the single-cell level was observed at the superior temporal gyrus. One of the neurons in our dataset showed selectivity to only two consonants (/SH/ and /JH/). Another neuron showed high selectivity to a specific diphthong (/OY/), as in ‘boy’, which is a combination of two vowels. The activity of such neurons suggest that single-cell activity can be highly selective in early auditory processing to specific sounds, to either consonants or to vowels, and that hierarchical processing may be observable at the single-cell level (e.g., from single phonemes to diphthongs).

The transition from modality-specific to amodal processing is a critical aspect in the transformation of sensory information to higher order linguistic features. Cross-modal invariance is a major step in hierarchical cognitive processing in general and in language processing in particular [46, 54, 55]. Although our design was not directed to the investigation of semantic features, previous research by Fried and Colleagues had discovered semantic concept cells in the human medial temporal lobe which are invariant to modality and can be reactivated during episodic memory [56, 57]. These cells have been recently shown to fire to pronouns [22]. The large spectrum of regional sampling in our study allowed for the demonstration of amodal responses in specific areas, the inferior frontal gyrus and frontal operculum as well as middle temporal gyrus. Several neurons in these areas were responsive to linguistic stimuli that were presented in both modalities. Although our sampling of single neurons in these regions was limited, as electrode placement was dictated solely by clinical considerations, the concomitant data from microwires and macro contacts allowed a more extensive sampling at various resolutions. Data obtained from these diverse methodologies, together with noninvasive electrophysiology and neuroimaging studies, hold considerable promise to shed light on the brain mechanisms underlying language function in humans.

The question of whether syntactic information is processed separately from semantic information in the human brain remains unresolved. Historically, a modular view was proposed, suggesting that syntactic information is processed in a dedicated system [e.g., 58–60]. Early neuropsychological studies supported this, identifying patients with impaired syntax but preserved semantics or vice versa [e.g., 61–64]. Brain-imaging studies, such as Dapretto and Bookheimer [65], initially seemed to confirm this modularity, localizing syntactic processing to Broca’s area. However, recent research, including the replication failure of Dapretto and Bookheimer [65], has led some researchers to challenge this view [2, 66] and may suggest a more integrated organization, with semantic and syntactic processing possibly involving intermingled neural circuits within partially overlapping cortical territories. Interestingly, in contrast to orthographic and phonological features, we have not identified, in the present dataset, any single cell selective to specific values of higher-level linguistic features, such as syntactic ones. It is possible that orthography and phonology follow a sparse code, with high redundancy, where single cells respond to interpretable features, whereas syntactic features are encoded in a more distributed manner. Such a high degree of overlap and feature intermingling has been observed in transformer models [67], which provide a relatively good fit to fMRI and MEG responses to written and spoken language in the areas where we found amodal responses [68–73]. Further research as to the neuronal basis of these features and the deciphering of sparse neuronal codes vs distributed population codes will require more extensive research with different methodologies subject to existing clinical opportunities for intracranial recordings in humans. Microwire recordings in the context of depth electrode implantation in epilepsy or other neurosurgical patients (sEEG) has the advantage of sampling of neurons in several brain regions simultaneously. Recent application of Neuropixel electrodes hold considerable promise of dense sampling with higher neuronal yield, albeit in a small cortical region, in brain tissue to be resected and for the foreseen future limited to intraoperative recordings of short duration [36].

## Methods and Materials

### Participants

We recorded single-unit activity from 21 patients at the UCLA neurosurgical centre with intractable epilepsy, who were implanted with Behnke-Fried electrodes [30, 31] as part of their clinical evaluation. All participants volunteered for the study by providing informed consent according to a protocol approved by the UCLA Medical Institutional Review Board (IRB).

### Stimuli

#### Design

The stimuli in the experiment were English sentences with length between two to five words. The sentences were created such that they contrasted various linguistic features by creating minimal comparisons. For example, to contrast the two values of grammatical number (singular vs. plural), the design contains minimal-pair contrasts, such as: “He smiles” vs. “They smile”; for gender “He smiles” vs. “She smiles”; for tense “He smiles” vs. “He smiled”, etc. (Figure 1A; see Table A1 for the list of contrasts). These contrasts along various grammatical dimensions allowed us to study, with a relatively small set of stimuli, neural activity in response to various linguistic features.

#### Auditory Stimuli

The auditory stimuli were recorded by a native Ametrican-English speaker, in a anechoic chamber. All stimuli were manually validated, pre-processed and the audio files were cut to include as minimal silence period at the beginning and end of the file. We used Montreal Forced Aligner (MFA^1^) to parse each auditory stimulus. MFA provides the onset times of each phone in the stimulus. Figure A1 illustrates the results for one of our stimuli, containing the sentence ‘He said that she smiled’. The results from MFA were further manually validated, to identify errors in the automatic parsing process. We used the results from the MFA to identify the actual sentence onset and offset, which we used in all analyses to replace the auditory-file onset and offset.

#### Paradigm

The stimuli were presented to the participants both (1) visually, using Rapid Serial Visual Presentation (RSVP) on a computer screen, with a word-presentation duration of 200ms and a stimulus onset asynchrony (SOA) of 500ms; and (2) auditorily, via standard computer speakers playing sentences recorded by a native American English speaker (Figure 1B). The experiment had six blocks in total, three visual and three auditory, which were presented intermittently, starting with a visual block. Each block contained the full set of stimuli, 152 sentences in total, repeated across blocks, each time in a random order. The participants were asked to perform a semantic task, and to press a button each time a word related to food was presented (which was the case in 5% of the stimuli; all removed from further analyses). To avoid eye movements, a fixation cross was presented at the beginning of each trial for a duration of 700ms.

### Data acquisition

Electrophysiological signals were recorded with Neuralynx and BlackRock devices, at high sampling rate for microwires (30,000-40,000Hz), required for extracting spiking data, and 2,000Hz for macro contacts.

### Preprocessing

Neural recordings from each patient were first manually examined to remove bad channels. The neural signal was then filtered to remove line noise at 60Hz, 120Hz, 180Hz and 240Hz (zero phase 2*^nd^* order Butterworth bandstop filters), and was normalized using robust scaling. Robust scaling removes the median and scales the data according to the quantile range. The neural signal was then clipped to be in the range of −5 to 5 interquatile ranges (IQR) from the median, where the IQR is the range between the 1st quartile (25th quantile) and the 3rd quartile (75th quantile). To extract high-gamma activity from microwires and macro contants, a frequency domain bandpass Hilbert transform (paired sigmoid flanks with half-width 1.5 Hz) was applied and the analytic amplitude was smoothed with a Gaussian kernel of width 25msec.

### Spike sorting

All units were semi-automatically sorted using Combinato [74]. When more than a single spike cluster was detected on a given microwire, the clusters were manually separated based on the shape of the spike, inter-spike interval distribution, and cross-correlation of the spike-times between clusters. Finally, all units were manually classified as single- or multi-unit based on the above factors and the presence of a refractory period. In this study, we included only units that were determined as single-unit.

### Encoding models

Temporal Receptive Fields (TRFs) of single cells and electrodes were estimated using generalized reverse correlation techniques [75], with regularized least-squares solutions [76]. These techniques were shown in the past to be effective for the modeling of intracranial recording from the human auditory cortex [77].

The TRFs included features from six feature groups (see Table A2): orthographic, phonological, lexical, positional, semantic and syntactic features. For each neuron and electrode, two TRFs were trained, one for each presentation modality (visual and auditory), with 92 and 94 features in total, respectively. To reduce model complexity and computation time, we reduced the sampling rate of the feature matrix to 20Hz. The feature matrix contained both binary and continuous features. For binary features, such as grammatical number, we used bipolar encoding, with ones denoting one of the possible values (e.g., ‘singular’ for ‘boy’), minus ones denoting the other value (e.g., ‘plural’ for ‘boys’) and zeros denoting either lack of a stimulus (e.g., silence periods) or irrelevant stimuli (e.g., grammatical number is irrelevant for ‘why’). For continuous features (e.g., log word frequency or sentence length), the stimuli were min-max scaled to the range [0, 1].

Each TRF was trained and evaluated with regularized least-squares techniques, using implementation from MNE-pyhton [78], in a cross-validation (CV) procedure with 20 folds. The Brain-Score of each TRF was defined as the mean Spearman correlation between the predicted and true neural activity across all 20 CV folds. This resulted in a single value per cell or electrode. To estimate the time-resolved Brain-Score, around, e.g., word or sentence onset, we also computed Spearman correlation between predicted and true neural activity per each time point. That is, we first predicted neural activity for all unseen trials, and then for each time point, we separately computed the Spearman correlation between the predicted and true neural activity. This resulted in a Brain-Score per each time point.

Feature and feature-group importance were computed using an ablation procedure. For each feature or feature-group of interest, we trained an ‘ablated’ TRF model with a feature matrix from which the feature-of-interest were removed.

## Acknowledgments

We are grateful to the participants for their involvement. We thank Gudamla Kalendar, Natalie Cherry, and Andreina Hampton for assisting in data collection and preprocessing. This work was supported by National Institute of Neurological Disorders and Stroke grant (R01NS084017, U01 grants NS108930 and NS123128) to IF. SD and YL were supported by INSERM, CEA, Collège de France, Université Paris Saclay, and an ERC advanced grant NeuroSyntax to SD.

## Author contributions

Conceptualization: YL, NF, SD, IF; Data analysis: YL, JK; Funding: IF, SD; Data acquisition: EM, AR, IF; Paradigm: YL, AT; Surgery: IF; Imaging: AR, IF. YL, IF wrote the manuscript. All authors provided critical review and commented on the manuscript.

## Competing interests

The authors declare no competing interests.

### Appendix A Stimuli

#### A.1 Contrasts and Feature Groups

**Table A1.** List of Main Contrasts in the Experiment.

| Contrast Name | Values | Example |
| --- | --- | --- |
| Speech Act | Declarative, Question | "He smiles.", "Who smiles?" |
| NP Type | Full noun, pronoun | "The boy smiles.", "He smiles." |
| Verb Type | Transitive, intransitive | "The girl kicked the ball.", "The girl smiles" |
| A-Movement | Unergative, unaccusative | "He smiled.", "He disappeared." |
| Grammatical Number | Singular, plural | "He smiles.", "They Smile." |
| Gender | Masculine, feminine | "He smiles.", "She smiles." |
| Tense | Past, present, future | "He sneezed.", "He sneezes.", "He will sneeze." |
| Question type | Subject, object | "Who smelled the flowers?", "What does the girl eat?" |
| Embedding | False, True | "The boy kissed the clown.", "He said that she smiled." |

For the encoding models, we defined six feature groups (Table A2). Each feature group is composed of several features from a specific aspect of language processing.

Categorical features are encoded using one-hot encoding. For example, part-of-speech in the lexcial group is encoded with a 5-dimensional vector. To encode, for instance, that a word is a noun, the first value of this vector is set to one and all others to zero.

**Table A2.** Feature Groups. Feature types used in each of the feature groups in the encoding TRF models.

|  | Feature Group | Features |
| --- | --- | --- |
| 1 | Orthographic | Letter identity, position |
| 2 | Phonological | Phonemes / phoneme features (voicing, ..) |
| 3 | Lexical | Word Frequency, Part-of-Speech |
| 4 | Positional | First word, last word, first phoneme, last phoneme |
| 5 | Semantic | GloVe (25 dimensions) |
| 6 | Syntactic | Grammatical number, gender, question |

#### A.2 Auditory Stimuli

**Fig. A1.**
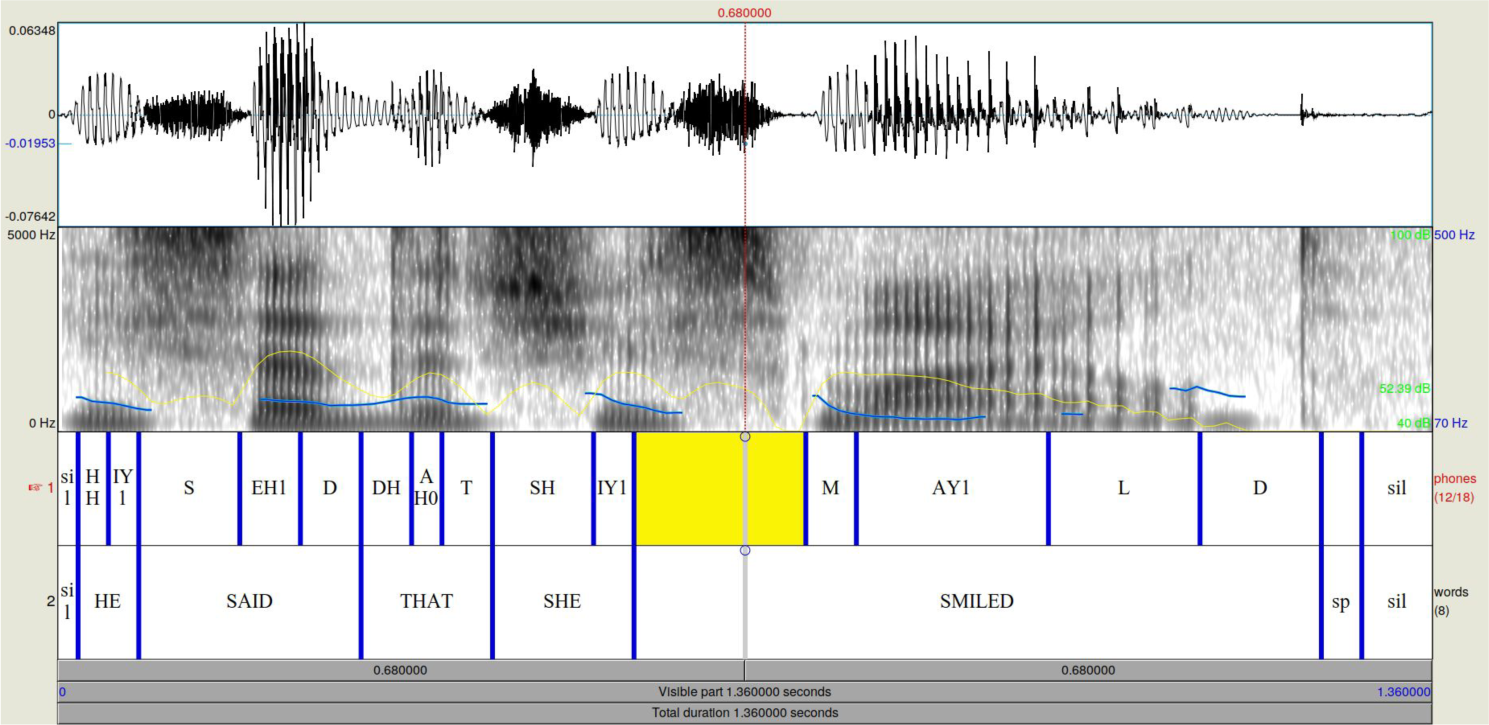
Auditory stimuli: An example of the waveform and spectrogram of one of the auditory stimuli in the experiment. The parsing to single phonemes and words is shown at the bottom two rows, based on Montreal Forced Alignment [79]. Light blue and yellow marking on the spectrogram refer to pitch and intensity, respectively.

### Appendix B Selective Single-Cell Responses in the Fusiform Gyrus

#### B.1 Modality Selectivity of Single Neurons in the Superior Temporal and Fusiform Gyrii

**Fig. B2.**
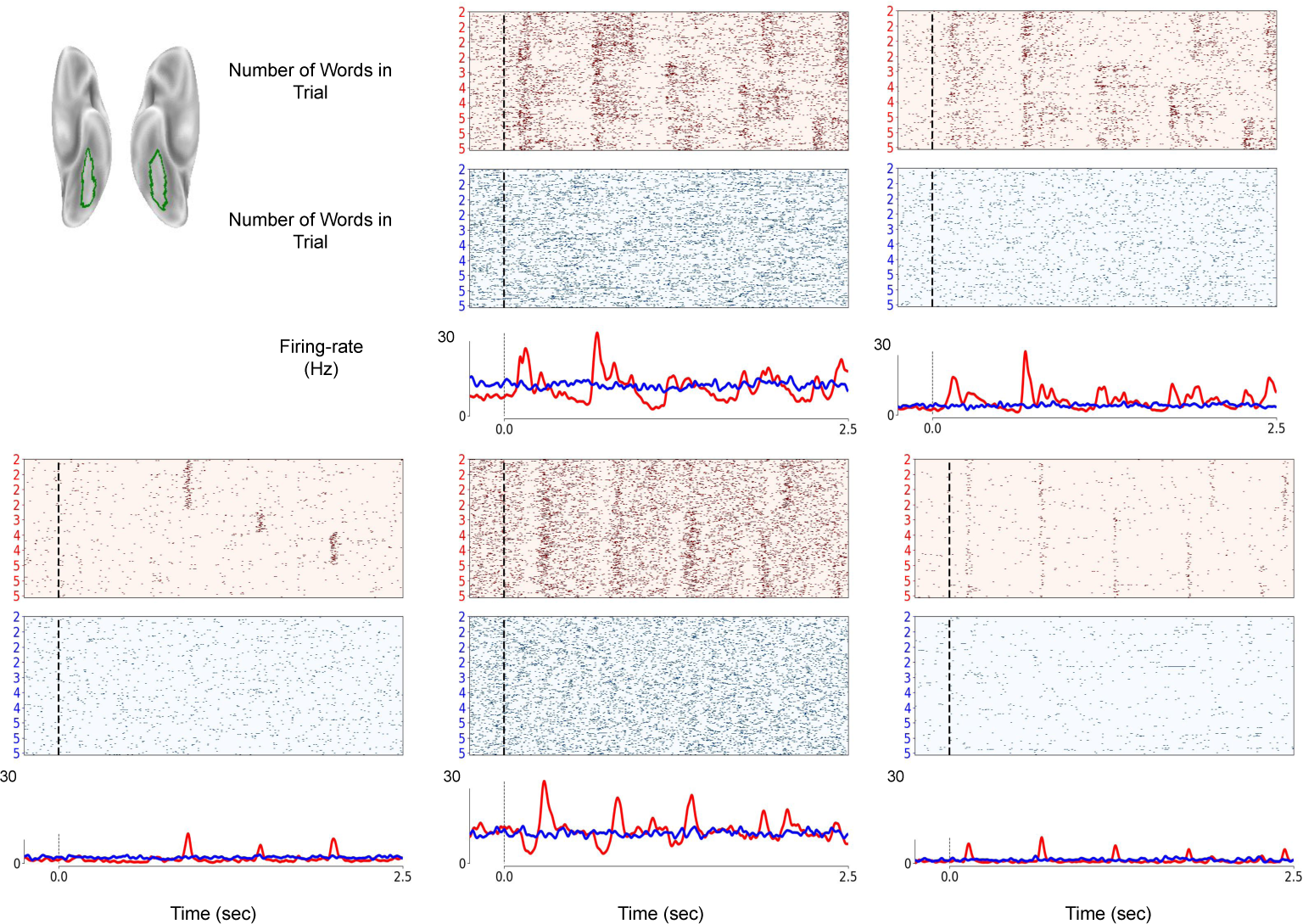
Modality Selectivity of fusiform neurons: Spiking activity from 5 neurons from the fusiform gyrus. All neurons show strong preference to the visual modality, remaining at baseline activity in response to audiory stimuli.

**Fig. B3.**
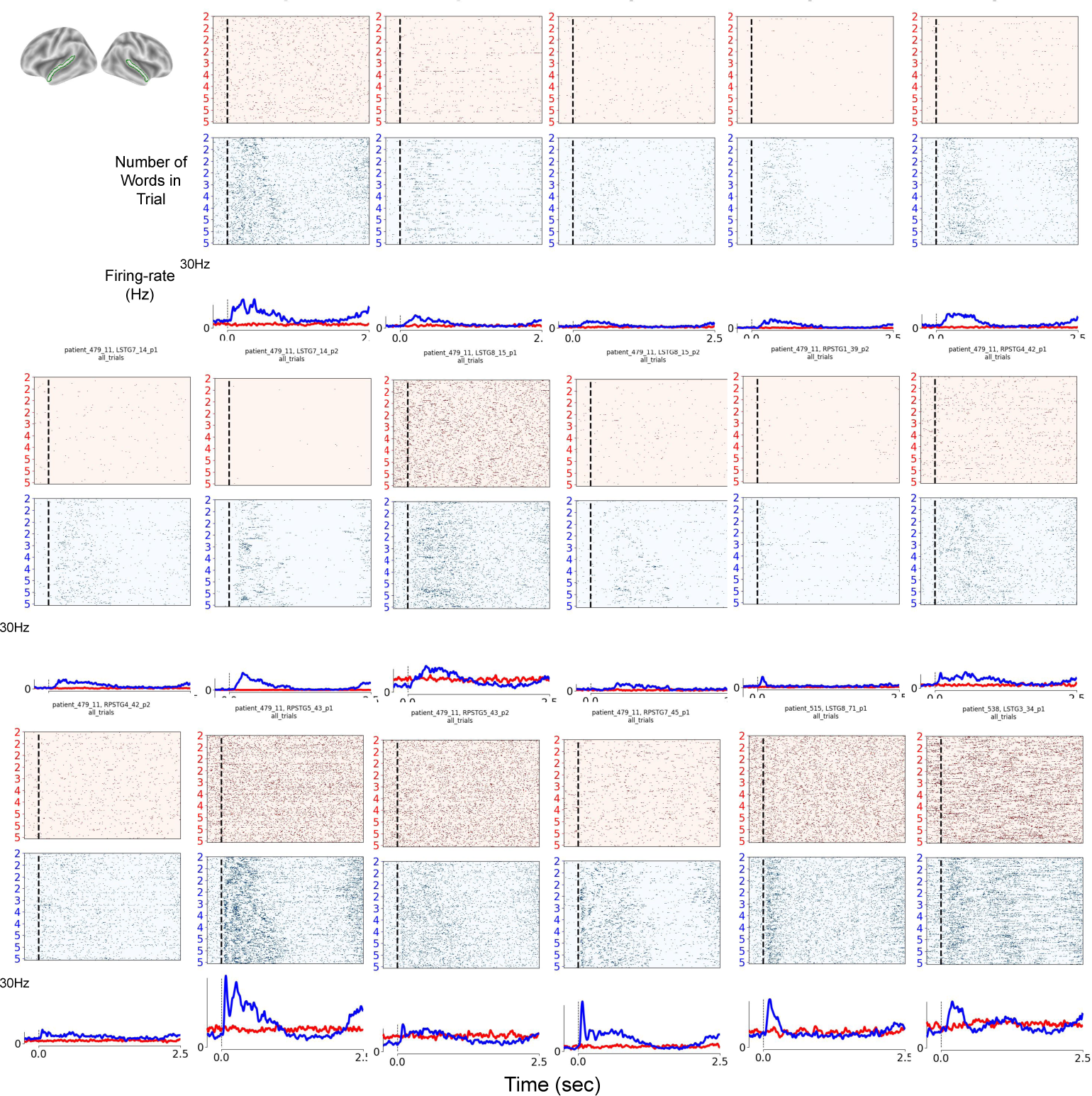
Modality Selectivity of STG neurons: Spiking activity from 17 example neurons from the Superior Temporal gyrus. All neurons show strong preference to the auditory modality, remaining at baseline activity in response to visual stimuli.

**Fig. B4.**
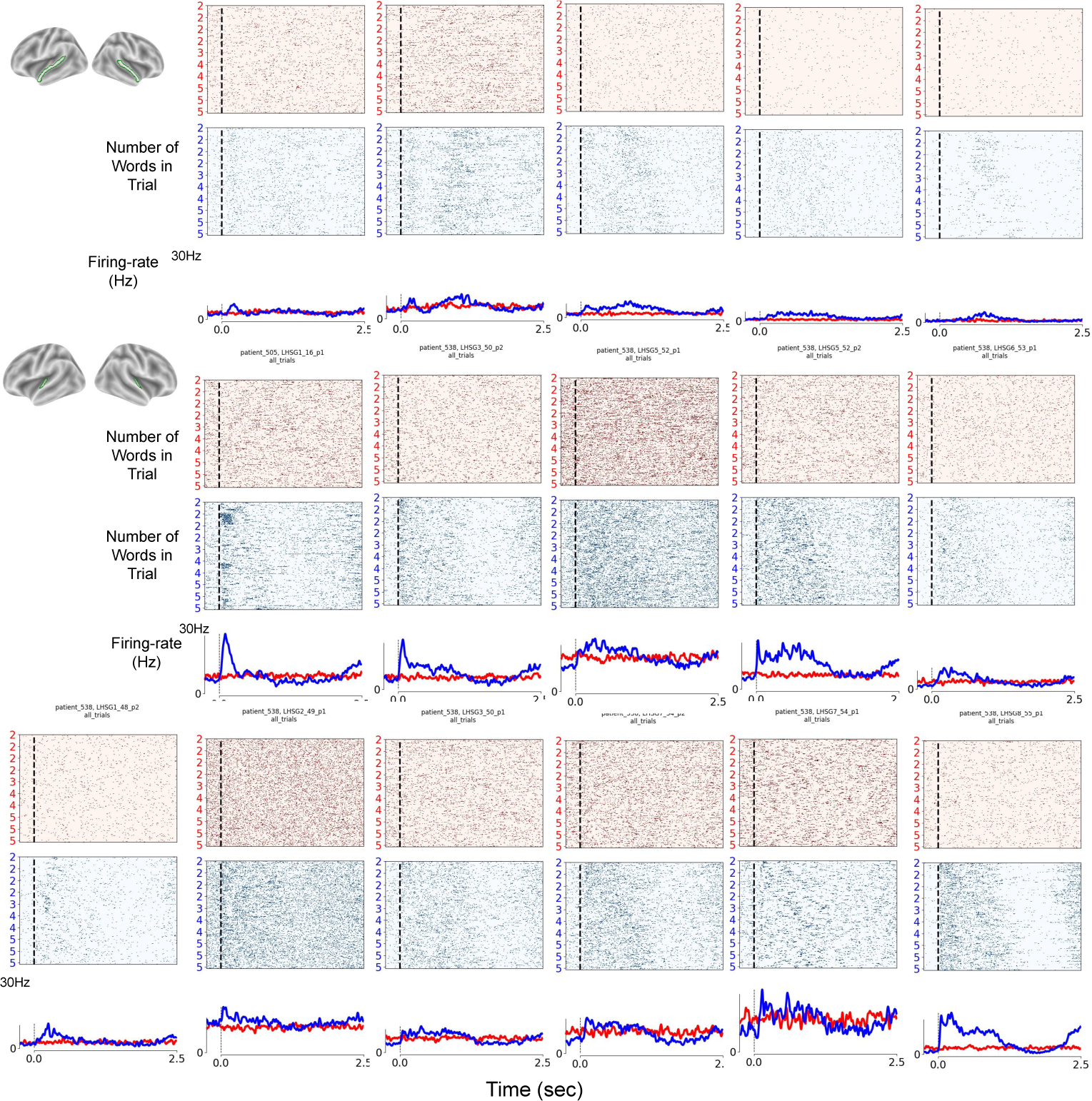
Modality Selectivity of STG and HSG neurons: Spiking activity from more 5 example neurons from the Superior Temporal gyrus and 11 from the Heschel Gyrus (HSG). All neurons show strong preference to the auditory modality, remaining at baseline activity in response to visual stimuli.

### Appendix C Selective Single-Cell Responses in the Superior Temporal Gyrus

**Fig. C5.**
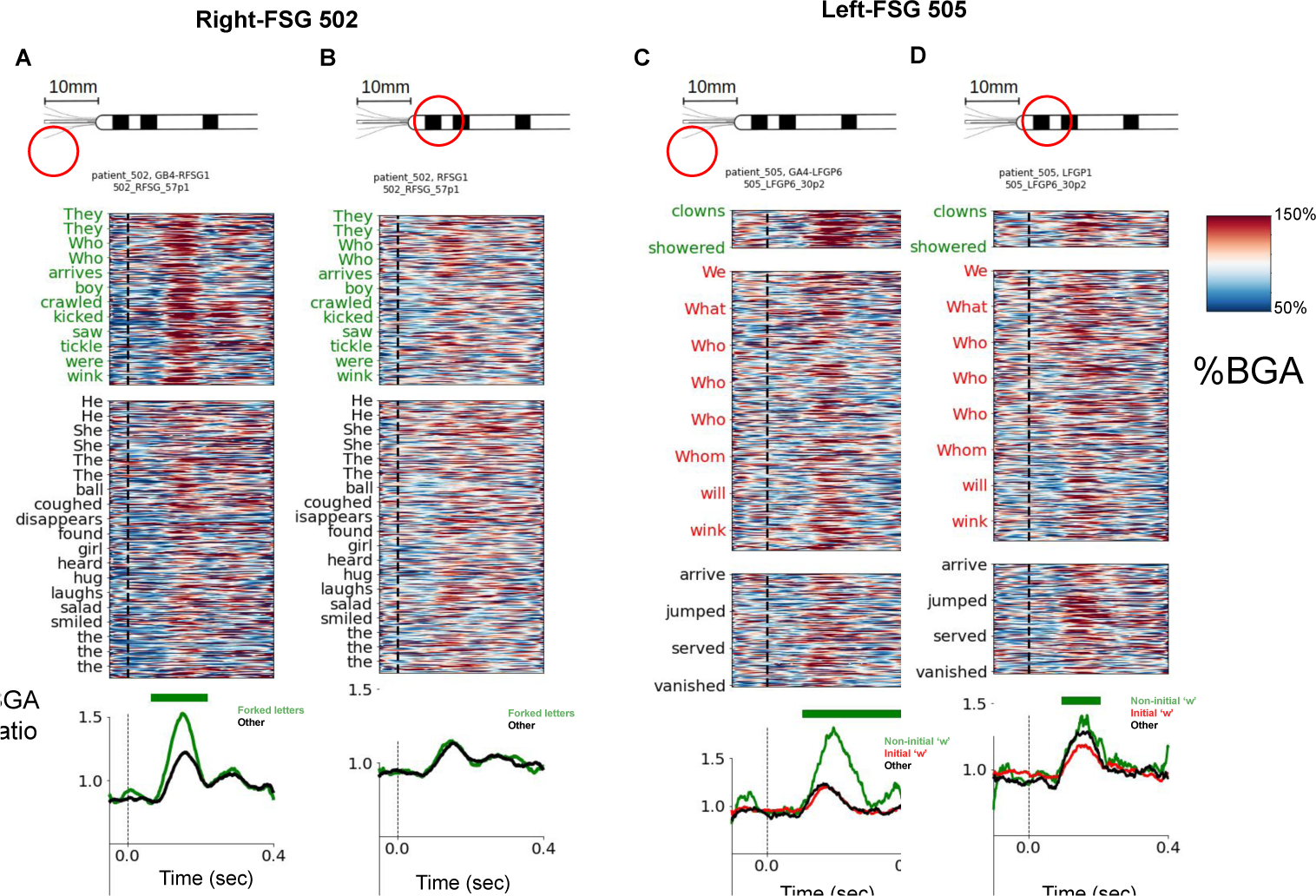
Micro and Macro-BGA Encoding of Orthographic Information during Visual Processing: (A) Micro broadband high-gamma activity (BGA) during the processing of words that contain forked letters (top panel) and those that do not (bottom). Micro BGA was extracted from the same microwire from which spiking activity was recorded in Figure 3A-D. Median BGA is shown at the bottom, separated for the two conditions. Time periods with significant separation (cluster-based permutation; *p − value <* 0.05 are marked with a green strip. (B) Same for macro BGA. (C+D) Micro and macro-BGA for the contrast shown in 3E-H. Significant separation is observed for micro-BGA, and to a lesser extent for macro BGA.

**Fig. C6.**
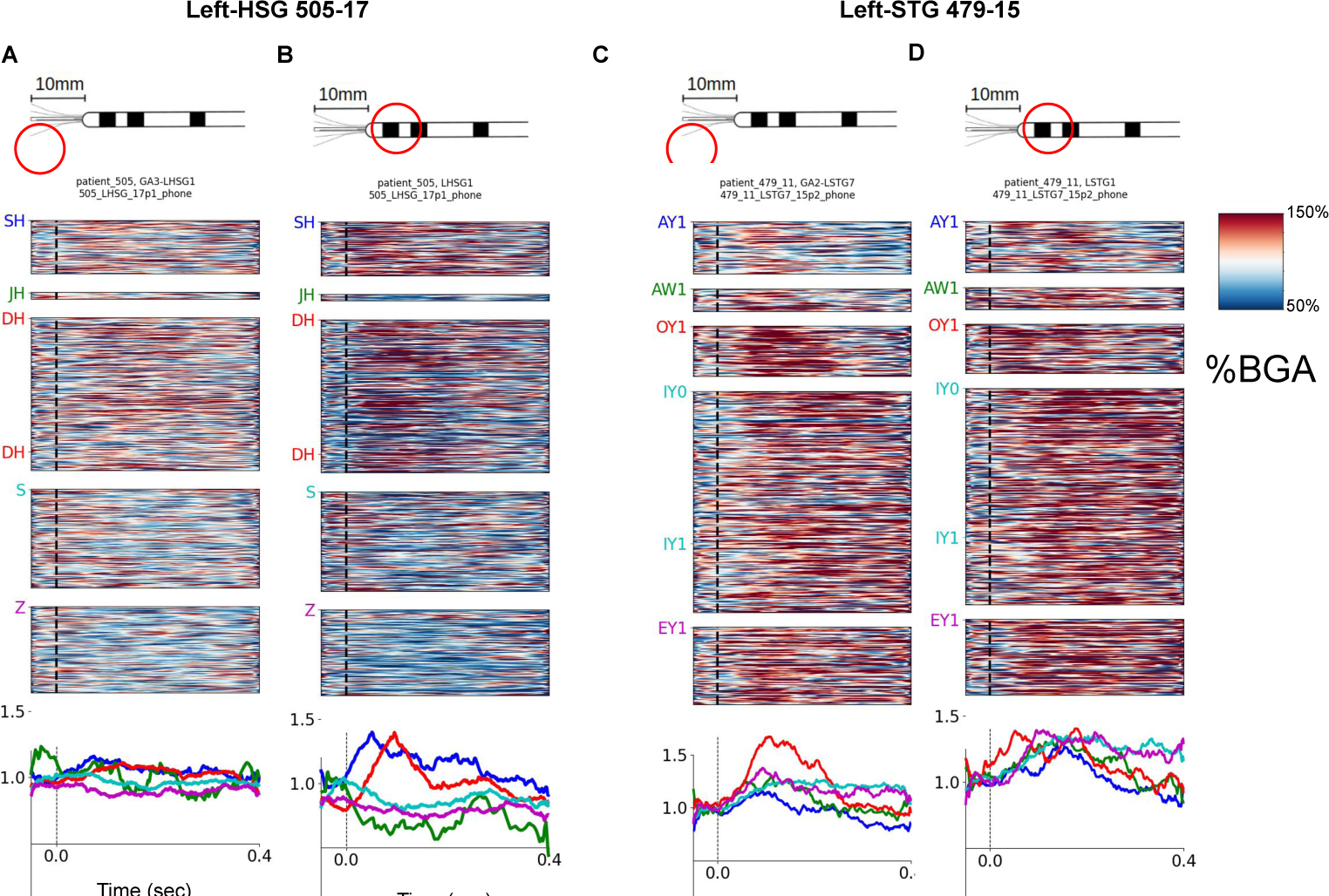
Micro and Macro-BGA Encoding of Phonological Information during Auditory Processing: (A) Micro broadband high-gamma activity (BGA) during the processing of words that contain hushing-sibilants (top panels) and those that do not. Micro BGA was extracted from the same microwire from which spiking activity was recorded in Figure 5A-D. Median BGA is shown at the bottom, separated for the two conditions. Time periods with significant separation (cluster-based permutation; *p −value <* 0.05 are marked with a green strip. (B) Same for macro BGA. (C+D) Micro and macro-BGA for the contrast shown in 5E-H. Significant separation is observed for micro-BGA but not for macro BGA.

### Appendix D Electrode Locations

**Fig. D7.**
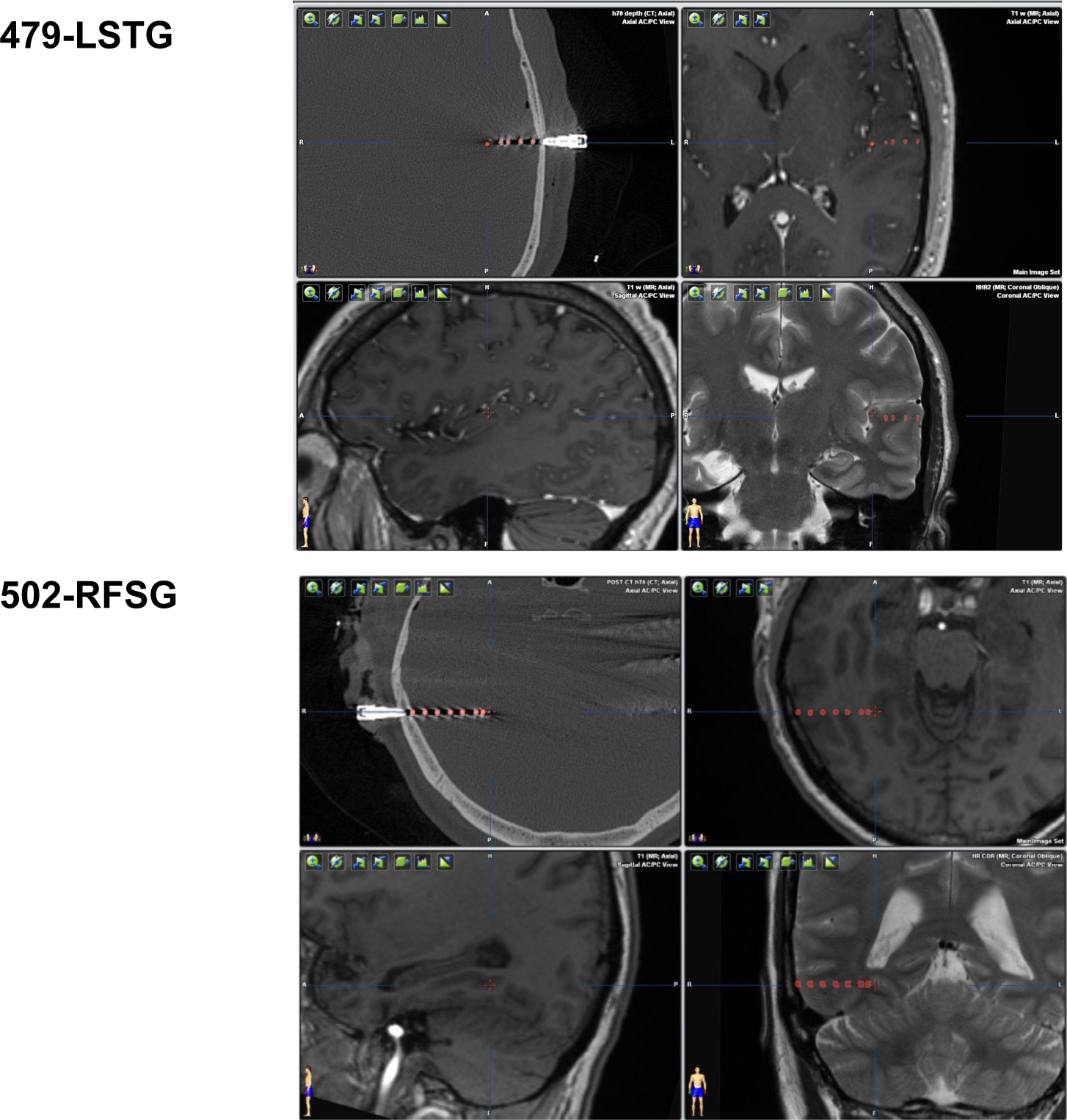
Electrode Localization.

**Fig. D8.**
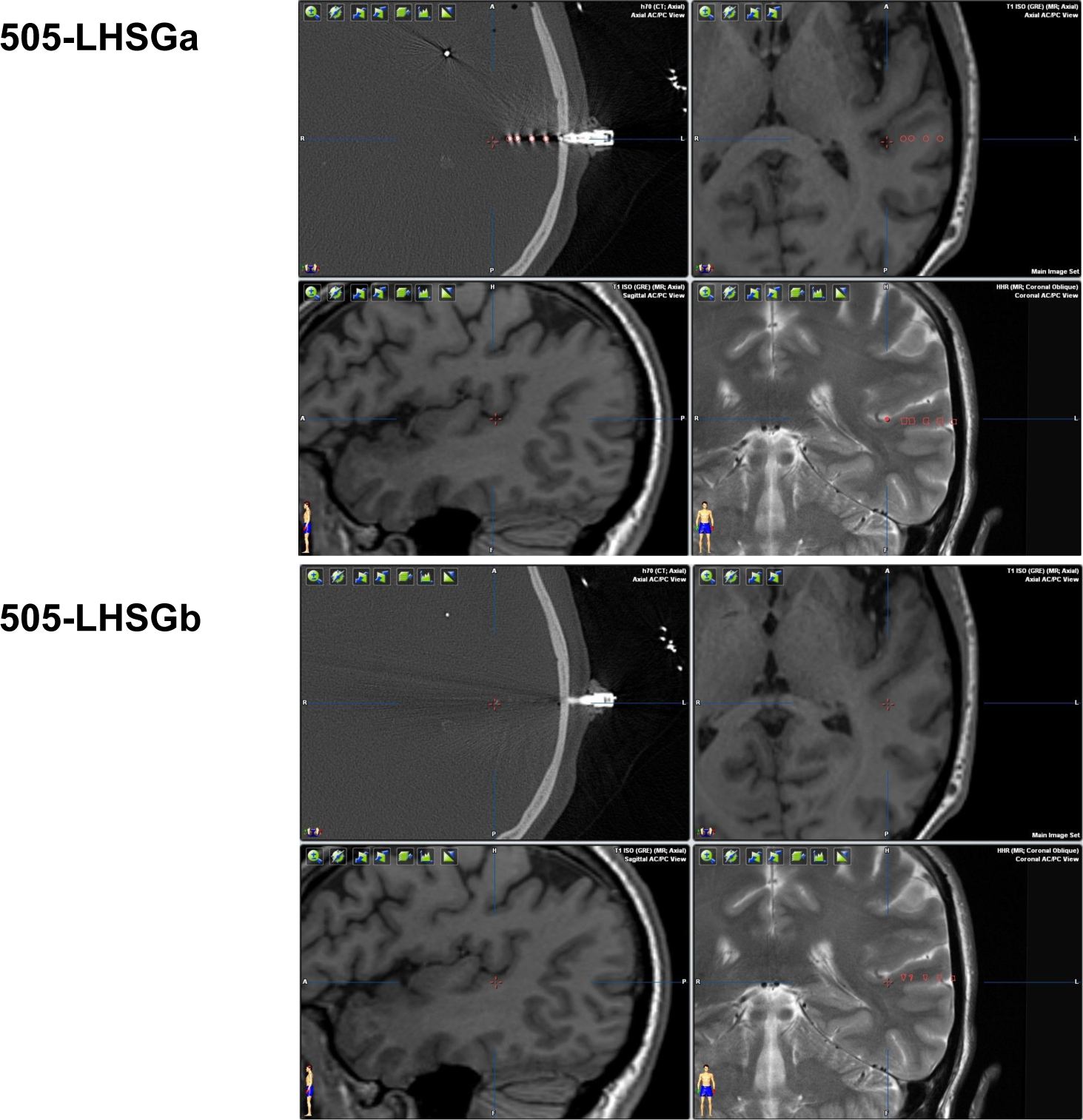
Electrode Localization.

**Fig. D9.**
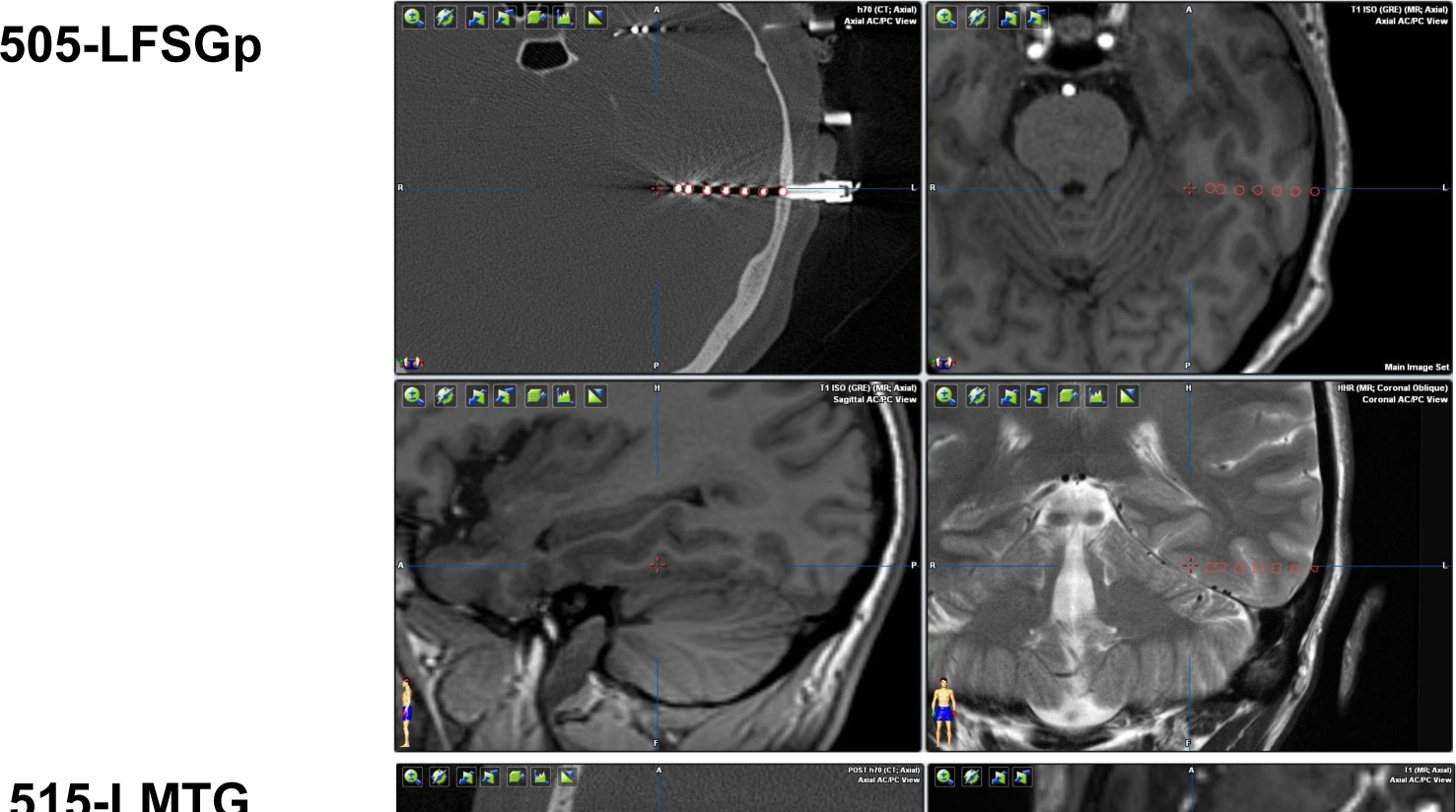
Electrode Localization.

**Fig. D10.**
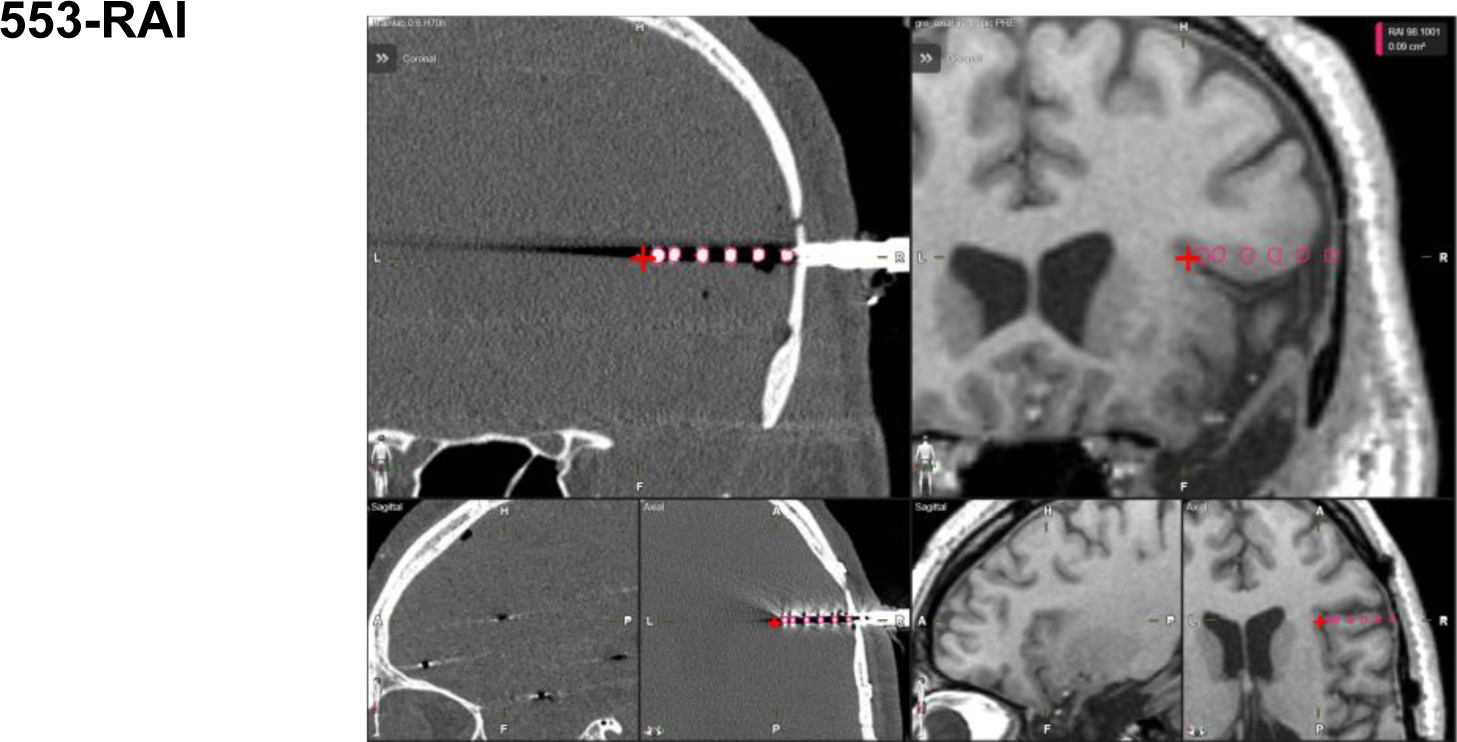
Electrode Localization.

### Appendix E Spike Sorting

We used Combinato [80] for spike sorting. One of the advantages of Combinato over other spike-sorting methods is that it optimizes the temperature of the supraparamagnetric-clustering algorithm [81]. Figure E11 shows the clustering results from Combinato for all units presented in the main text.

**Fig. E11.**
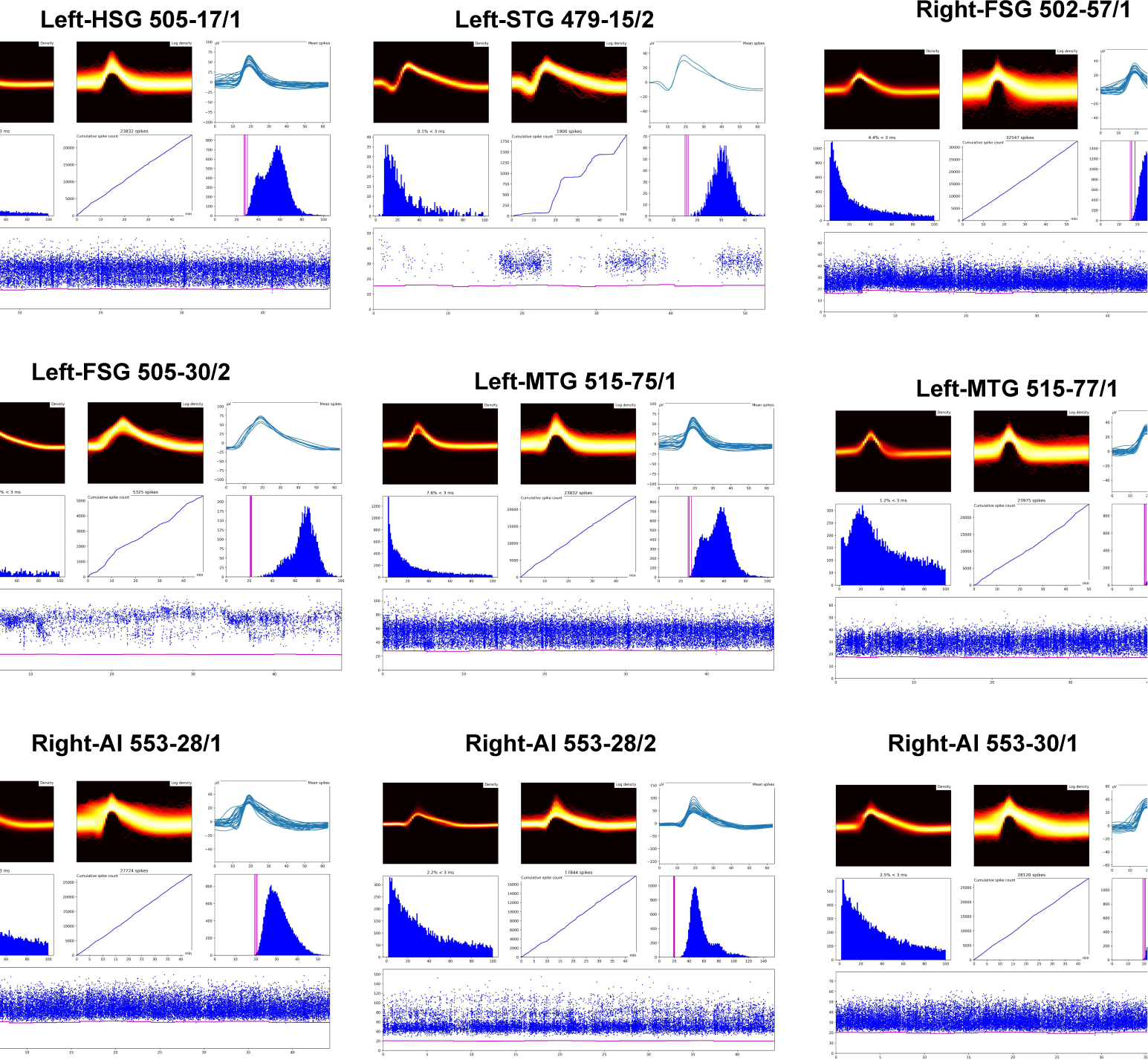
Spike Sorting: For each unit, the results of the spike sorting are summarized in: (1) the profile of the cluster (top row, including in log scale), (2) the inter-spike-interval distribution (middle-left panel), (3) the cumulative spike count (center), (4) the distribution of the spike amplitude (middle-right panel), and (5) amplitude of all spikes across the entire experiment (bottom row).

### Appendix F Neural Encoding of Word Length

**Fig. F12.**
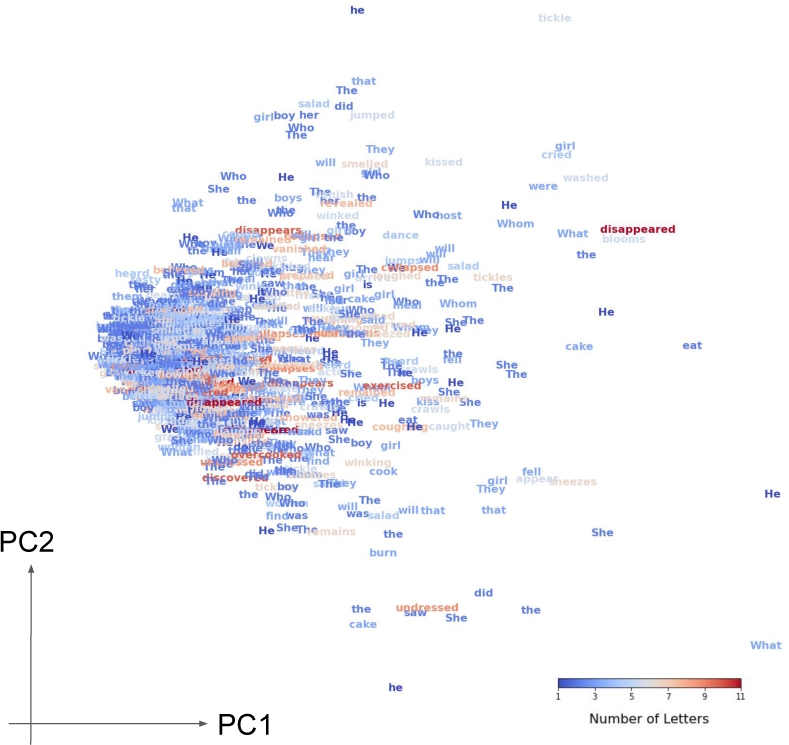
Neural Encoding of Word Length (micro BGA): (A) Principal Component Analysis (PCA) of micro BGA activity from the 5 fusiform neurons with the highest Brain-Score. The neural response of each neuron was represented along 5 consecutive 100ms time bins between 0.1-0.6sec after word onset. The population response to different words is projected onto the two main PCs (39.7% and 9.9% of the variance). Words are colored by their length.

### Appendix G Feature Importance

**Fig. G13.**
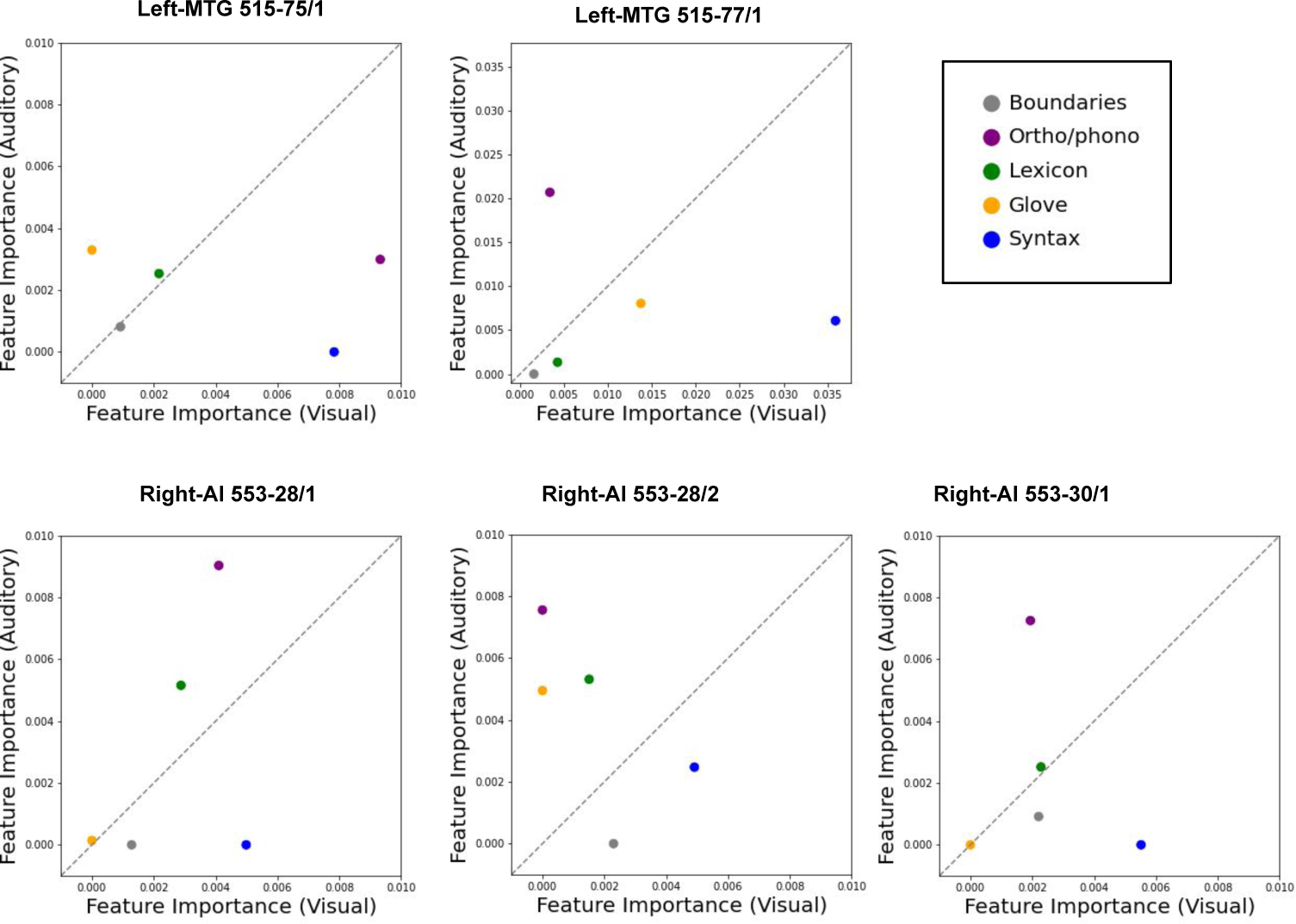
Feature Importances for Amodal Neurons: For each amodal unit in Figure 6, the corresponding scatter contrasts group feature importances (FI) for the two blocks (visual and auditory). Each point in a scatter corresponds to a feature group (see legend for color code). Points on the diagonal represent FIs that are similar across the two blocks.

## Footnotes

1 https://montreal-forced-aligner.readthedocs.io/en/latest/index.html

## Notes

### Competing Interest Statement

The authors have declared no competing interest.

### Summary of Updates

Correction of author details. The first and last names of the first author were reversed in the original submission.

